# Inferring gene-pathway associations from consolidated transcriptome datasets: an interactive gene network explorer for *Tetrahymena thermophila*

**DOI:** 10.1101/2024.12.12.627356

**Authors:** Michael A. Bertagna, Lydia J. Bright, Fei Ye, Yu-Yang Jiang, Debolina Sarkar, Ajay Pradhan, Santosh Kumar, Shan Gao, Aaron P. Turkewitz, Lev M. Z. Tsypin

## Abstract

Although an established model organism*, Tetrahymena thermophila* remains comparatively inaccessible to high throughput screens, and alternative bioinformatic approaches still rely on unconnected datasets and outdated algorithms. Here, we report a new approach to consolidating RNA-seq and microarray data based on a systematic exploration of parameters and computational controls, enabling us to infer functional gene associations from their co-expression patterns. To illustrate the power of this approach, we took advantage of new data regarding a previously studied pathway, the biogenesis of a secretory organelle called the mucocyst. Our untargeted clustering approach recovered over 80% of the genes that were previously verified to play a role in mucocyst biogenesis. Furthermore, we tested four new genes that we predicted to be mucocyst-associated based on their co-expression and found that knocking out each of them results in mucocyst secretion defects. We also found that our approach succeeds in clustering genes associated with several other cellular pathways that we evaluated based on prior literature. We present the *Tetrahymena* Gene Network Explorer (TGNE) as an interactive tool for genetic hypothesis generation and functional annotation in this organism and as a framework for building similar tools for other systems.

**Key Points:**

- ur approach integrates nearly 20-year-old microarray and contemporary RNA-seq datasets.
- rigorously compare co-expression clustering parametrization by way of computational controls.
- Co-expression clustering identifies known and novel functionally associated genes in *Tetrahymena*.

**Graphical Abstract:** 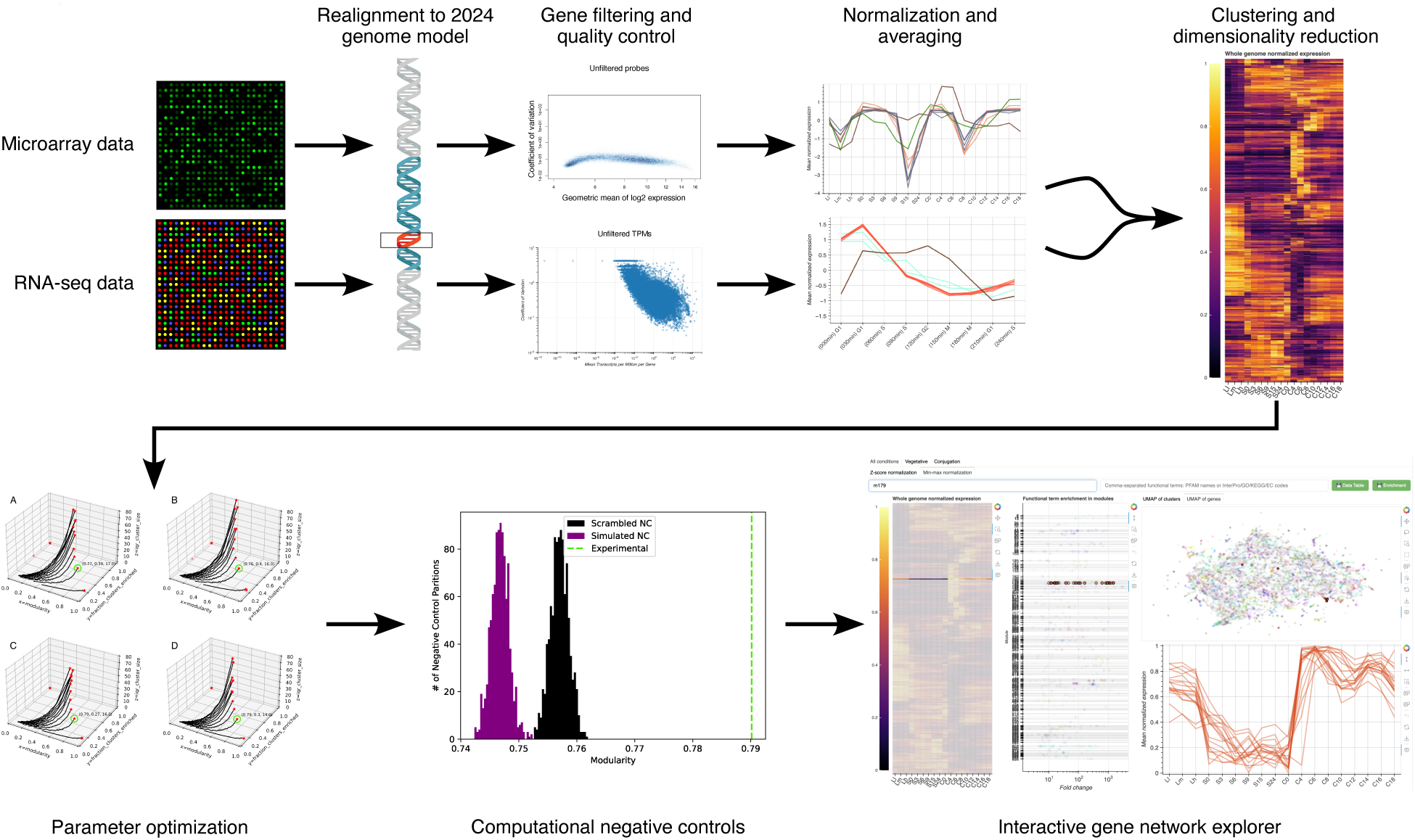

## Introduction

Gene co-expression, particularly in response to an experimental perturbation, has long been used as evidence for the functional association of genes that are otherwise uncharacterized (1). The transcriptome is an intermediary between genotype and phenotype, and transcriptomics is often cheaper, faster, and higher throughput than using biochemistry or genetic engineering to functionally associate a given gene with a biological pathway or process (2). The number of transcriptomic datasets has grown dramatically over the past two decades, raising deep questions such as: how well do co-expression patterns translate from one set of experimental conditions to another? How many cellular processes are driven by genetic co-expression, and does this change under different growth or environmental conditions? These questions point to the importance and challenge of using the wealth of publicly available data to pursue new hypotheses, rather than treating whole-transcriptome experiments as either purely descriptive or one-and-done assays to study a single organism- or condition-specific problem. Answering these questions requires appropriate model systems and principled approaches.

The ciliate *Tetrahymena thermophila* is a unicellular eukaryote that has featured in groundbreaking discoveries regarding programmed genome rearrangements, telomeres/telomerase, and cytoskeletal motor proteins (3). However, some features of *T. thermophila* present challenges to its use in uncovering new biology broadly. Ciliates are over a billion years diverged from better studied organisms such as fungi and animals (4), an evolutionary distance that frequently creates obstacles to identifying gene orthologs and thereby inferring conserved functions. Moreover, interesting novel mechanisms may have arisen over that large evolutionary distance, such as the recently discovered unique secretory apparatus that ciliates share only with the related apicomplexans and dinoflagellates (5). One way to address these issues would be a forward genetic approach, using random mutagenesis to identify phenotypes of interest and then associate them with causative mutations (6–8). However, ciliate nuclear organization makes it challenging to undertake high throughput forward genetic approaches in these organisms (9). Due to all these factors, high throughput bioinformatic studies offer a potential breakthrough for interrogating both the evolutionarily conserved and novel biology in *T. thermophila*. Previous research has indicated that protein expression in *T. thermophila* tends to be regulated on the level of transcription as opposed to transcript degradation or translation rates, which is in line with observations in yeast (10, 11). Thus, gene co-expression studies promise to be informative for the analysis of gene functions.

*T. thermophila* has distinct vegetative and sexual life stages. Consequently, many genes are tuned for differential expression during stages of vegetative growth/mitosis or conjugation/meiosis, previously explored in microarray-based experiments and co-expression analyses (12–15). Strikingly, we found that many characterized genes involved in the biosynthesis and secretion of a particular secretory organelle, called the mucocyst, are co-expressed across growth, starvation, and conjugation (16–18). This allowed us to subsequently identify multiple other co-expressed genes. A large subset of these were then verified as involved in mucocyst biogenesis or secretion (16–18). This success led us to develop a tool we called the Co-regulation Data Harvester (CDH), which scraped the available co-expression data for *T. thermophila* and performed reciprocal-best-BLAST queries to indicate potential functional annotations based on orthologous genes in other organisms (19). This tool also allowed us to identify candidates for genes involved in the secretion of homologous organelles in the apicomplexan *Toxoplasma gondii*, which were then experimentally verified (20).

However, the CDH became obsolete as new algorithms for identifying gene co-expression, as well as new databases for orthology searches, became available after our publication (21–24). Additionally, extensive new revisions of the *T. thermophila* genome model were published, as well as an RNA-seq dataset from cell cycle-synchronized cultures (25–27). These developments prompted us to align the original microarray data with the newest genome model, while also bridging insights from the different transcriptomic datasets, to develop a stable, accessible tool for the research community. Here, we report the *Tetrahymena* Gene Network Explorer (TGNE), an interactive tool for revealing co-expression patterns, by taking advantage of the gene annotations and expression data that have only recently become available (21, 22, 28). One important aspect of the expanded datasets is that the microarray and RNA-seq expression profiles are independent from each other, the former covering bulk growth, starvation, and sexual conjugation, and the latter covering a synchronized mitotic cell cycle. Using the TGNE, we found that co-expression of many mucocyst-related genes is a feature of both datasets. We further found that these genes are also upregulated during experimentally induced mucocyst biosynthesis, implying that their co-expression in “untargeted” experiments reflects functional association. This demonstrates that the TGNE can be used to generate experimentally verifiable hypotheses and provides a direct insight into the dynamics of functionally associated genes in *T. thermophila*. A similar pattern emerges from TGNE analysis with regard to other cellular processes in *T. thermophila*, such as regulation of histone, ribosomal, and proteasomal subunits.

Beyond drawing insights specific to *T. thermophila* biology, our approach to developing the TGNE provides a framework for revitalizing microarray data and integrating it with RNA-seq results. We leverage computational negative controls to support our choice of (dis)similarity metric for co-expression profiles and our optimization of parameters for partitioning the profiles into clusters. We also compare different normalization strategies to show the degree to which gene expression pattern shape and magnitude differently affect the clustering results. Approaches to computational negative controls, distance metrics, normalization, and clustering algorithms have all been detailed in prior work (1, 24, 29–32). However, to our knowledge, these strategies have not been previously brought together to unite bioinformatic insights from different experiments. Our results indicate that there are more testable hypotheses to be found and more insights to glean from the troves of publicly available bioinformatic data, even for evolutionary distant and experimentally challenging organisms.

## Materials and Methods

### Code and data availability

All code and data necessary to reproduce our analysis is available through Zenodo (https://doi.org/10.5281/zenodo.14353373) and figshare (https://doi.org/10.6084/m9.figshare.28022501). Our new microarray dataset for mucocyst regeneration after secretion is available on the NCBI Gene Expression Omnibus: https://www.ncbi.nlm.nih.gov/geo/query/acc.cgi?acc=GSE276404.

### Microarray co-expression analysis

The microarray data analyzed in this study were sourced from the NCBI Gene Expression Omnibus (GEO; https://www.ncbi.nlm.nih.gov/geo/) under accession numbers GSE11300, GSE26650, GSE26384, and GSE26385. Experimental probes were aligned to the June 28, 2024 release of the *Tetrahymena thermophila* genome model CDS using HISAT2 with default parameters (27, 33). In the NimbleGen Design File (NDF), the SEQ_ID of each singly aligned probe was replaced with the TTHERM_ID of the corresponding sequence. Probes that did not align to a single gene coding sequence were discarded with the exception of RANDOM probes. All raw microarray data files were converted to XYS format. All XYS files were compiled to create an expression set and RMA normalized with oligo and pdInfoBuilder in R (34, 35).

### Microarray chip quality control

Chips were removed if they met the following three criteria: (i) had a 25th percentile NUSE (normalized unscaled standard error) > 1, (ii) had a relatively large NUSE interquartile range, and (iii) had expression intensity autocorrelation on reconstructed pseudo-images of the original chips (36). After this quality control, if there remained only one replicate for a given time point, it was also excluded from the analysis. The microarray chips were hierarchically clustered with hclust in R to observe any clustering biases. All microarray chips from GEO Accession GSE26385 were removed, as they clustered away from other replicates for their respective conditions and were collected by a specific individual, which is indicative of a batch effect (13).

### Microarray gene filtering

Genes were filtered to remove ones that had too low expression or variance to be informative in the analysis. All genes were subjected to two filters: one based on the distribution of their respective expression statistics and one based on likelihood of differential expression. The first filter required that:

1. The gene’s geometric mean expression was greater than or equal to the 25th percentile of the geometric means of expression for all genes; OR,
2. The gene’s geometric coefficient of variation of expression was greater than or equal to the median geometric coefficient of variation for all genes; OR,
3. The gene’s maximum fold-change of expression was greater than or equal to the median maximum fold-change of expression for all genes; OR,
4. The gene’s ratio of its median absolute deviation to its median expression was greater than or equal to the median ratio for all genes.

To identify genes that have robust differential expression patterns that may have been lost to the above filter, we used MaSigPro parametrized to a false discovery rate q = 0.001 by the Benjamini-Hochberg correction for multiple hypothesis testing (37). The genes identified by MaSigPro were added to the ones that passed the first filter, and this total gene set was used for subsequent analysis.

### RNA-seq analysis

The sequencing data used in this study was downloaded from the Sequence Read Archive (SRA; https://www.ncbi.nlm.nih.gov/sra/) under the BioProject PRJNA861835. Adapter sequences and low-quality reads were removed with Trimmomatic (38) with default parameters. The quality of the reads in each sample before and after trimming was accessed with FastQC (39) and MultiQC (40). The transcript abundance for each gene was computed with Kallisto (41) using the trimmed reads and the *T. thermophila* genome model CDS. Transcripts per million (TPM) and counts per million (CPM) were computed from transcript abundance and the effective length of each transcript.

### RNA-seq gene filtering

Jaccard filtering was applied to the RNA-seq gene expression data to remove genes with noisy and unreplicated expression patterns (42). We determined the maximum Jaccard similarity index between replicate gene measurements to be 0.9422, which corresponded to a CPM of 0.0802. Only genes with a maximum CPM measurement above 0.0802 were kept for the subsequent analyses. After filtering, the TPM values for expression were used to compute co-expression clustering.

### Orthology-based annotation

eggNOG-mapper v2.1.12 was applied to the *T. thermophila* genome model protein sequences to mine annotations of orthologs using the following parameters: the HMMER database with 2759 as the taxID, tax scope constrained to Eukaryota, 2759 selected for the target taxa, report orthologs enabled, non-electronic GO terms only, the HMM database, and no PFAM realignment (21, 22, 43). The exact command was: -m hmmer -d 2759 --no_annot --tax_scope Eukaryota --target_taxa 2759 --report_orthologs --report_no_hits --go_evidence non-electronic --pfam_realign none --dbtype hmmdb. Interproscan 5.68-100.0 was applied to the *T. thermophila* genome model protein sequences to mine annotations of orthologs using the default parameters (28, 44).

### Enrichment analysis

The modified two-tailed Fisher’s Exact Test with a Bonferroni correction for multiple hypothesis testing, as implemented in the DAVID database, was used to determine if any GO, COG, EC, KEGG_ko, PFAM, and/or InterPro annotation terms were enriched in each cluster relative to the genome background (45, 46).

### Clustering

The raw microarray and RNA-seq expression datasets were preprocessed and clustered using the same approach. Two different preprocessing pipelines were applied to each dataset: (i) each gene expression profile was log-transformed elementwise and subsequently z-score normalized; (ii) each gene expression profile was min-max normalized. All four preprocessed datasets were subject to the same clustering pipeline. The arithmetic mean of the normalized expression values was computed for each gene across replicates at each time point. A high-dimensional Manhattan distance matrix was precomputed with scikit-learn (47). The nearest neighbors for each gene expression profile in the high-dimensional space were computed using a modified scikit-learn function. By default, scikit-learn does not include a point as its own nearest neighbor. The scikit-learn function was used to compute the eleven nearest neighbors, and the point itself was added manually to the set of nearest neighbors and distances to complete the set of twelve. A graph of the high-dimensional space was built with umap-learn (23) with a varying number of nearest neighbors. Genes were clustered via community detection of networks with leidenalg (24) using the Constant Potts Model (48) quality function with a varying linear resolution parameter.

### Parameter optimization and partition quality validation

Partitions were computed with a varying number of nearest neighbors for UMAP graph generation (range=[2, 12], step=1) and Leiden linear resolution parameters (range=[0, 1], step=0.005). The modularity of each partition was computed using the graph and the clusters generated by the partition with networkx (49). The number of enriched clusters was counted, and the fraction of enriched clusters was computed as the number of enriched clusters divided by the total number of clusters in the partition. The interquartile range among all of the cluster sizes was computed for each partition. Pareto-efficient partitions were computed in a three-dimensional space defined by the fraction of enriched clusters, modularity, and the interquartile range of cluster sizes. For a partition to be considered optimal, we required a modularity greater than 0.7 and an interquartile range of cluster sizes greater than 10. Computational negative control partitions were used to assess the statistical significance of each of the four optimal partitions. Scrambled negative control partitions were generated by randomly swapping raw expression values within each gene’s expression profile before preprocessing and clustering. Simulated negative control partitions were generated by creating a uniformly distributed Latin hypercube with SciPy (50) with the same dimensionality as the dataset. The hypercube values were then scaled to match the range of values within the dataset and used as raw input for preprocessing and clustering. 1000 computational negative control partitions of each type were computed with the optimal parameterizations for each of the microarray and RNA-seq datasets. A two-tailed one-sample t-test was used to assess the difference between the modularity of each optimal partition and each of the corresponding negative control modularity distributions.

### Differential expression analysis of induced mucocyst replacement dataset

To distinguish between genes upregulated due to demands of mucocyst synthesis vs. genes upregulated in response to the exocytic stimulus *per se*, we exploited a mutant cell line, MN173, that does not secrete its mucocysts upon stimulation (51). Using biological triplicates, total RNA was isolated from wildtype cells (strain CU428) prior to stimulation of exocytosis, and then 60 minutes post stimulation. In parallel, cells from the MN173 mutant line were treated and processed equivalently. Cells were grown in 1% proteose peptone, 0.2% dextrose, 0.1% yeast extract, 0.003% ferric EDTA. Cells were grown to 150, 000-200, 000 cells/ml and pelleted in 50ml conical tubes for 45 sec at 800xg. They were washed once and suspended in DMC (0.1 mM Na2HPO4, 0.1mM NaH2PO4, 0.65mM CaCls, 0.1mM MgCl2, pH7.1) for 16 h at room temperature with shaking. 50 ml aliquots were stimulated by pelleting as above, and resuspension in 5 mL. 2% Alcian Blue was added to 0.05% and the tube contents were mixed by inversion, and then diluted immediately by addition of 45 mL of 0.25% proteose peptone, 0.5mM CaCl2. Cells were then washed once in DMC and resuspended for recovery in DMC at room temperature with shaking. Pelleted cells were lysed with 5M guanidinium thiocyanate, 10mM EDTA, 50mM Tris-HCl pH7.5, 8mM 2-mercaptoethanol. RNA was precipitated with 7 volumes cold 4M LiCl, and the pellet was washed once with 3M LiCl and suspended in 0.5% N-lauroyl-sarkosine, 1mM EDTA, 10m Tris pH7.5. RNA was then phenol/chloroform extracted and ethanol precipitated.

The cDNA synthesis and Cy 3 labeling were done by Roche NimbleGen Systems, as described in (52). Hybridization and staining of arrays were carried out by Roche NimbleGen Systems as described in (53). Arrays were scanned by Roche NimbleGen using a GenePix 4000B (Molecular Devices, Sunnyvale, CA) and the data were extracted using NimbleScan software. Array normalization was performed using the quantile normalization method (54). Normalized expression values for the individual probes were used to obtain the expression values for a given open reading frame by using the multiarray average (RMA) procedure (55). Data were analyzed based on the RMA-processed expression values.

The microarray data from the mucocyst replacement experiments were processed the same way as the untargeted co-expression microarray dataset up to and including the RMA normalization step. The RMA normalized expression set was analyzed with limma (56) to determine the differential expression data for each gene over one hour in the MN173 mutant relative to the wild type *T. thermophila*. Genes which had a fold-change of greater than 1.5x, a Benjamini and Hochberg’s adjusted false discovery rate (q-value) lower than 0.01, and a B-statistic greater than 1 (i.e., a Bayesian posterior probability greater than a 73.1% chance of being differentially expressed) were classified as differentially expressed genes.

### Cross-validation of untargeted co-expression analyses against targeted mucocyst replacement experiment

We identified co-expression clusters that were statistically significantly enriched for the 33 genes that are experimentally validated to be involved in mucocyst biogenesis (Supplementary File 1) in both the microarray and RNA-seq datasets. This was done by a Fisher’s Exact Test relative to the background genome as in (45). For each respective normalization framework, these sets of genes and the set of genes that are upregulated during mucocyst replacement were then compared to determine their mutual agreement. A Venn diagram was generated to display the intersections of the three sets. To assess whether the intersections between these sets are more likely to include experimentally validated genes than the genes excluded from the intersections, we again employed a Fisher’s Exact Test, this time looking at the background of the union of the three sets, rather than the entire genome.

### Macronuclear knockouts of candidate genes

Genes of interest (TTHERM_00141040, TTHERM_00193465, TTHERM_01213910, TTHERM_00047330, TTHERM_00317390, TTHERM_00283800, TTHERM_00241790, TTHERM_01332070, TTHERM_00059370, and TTHERM_00227750) were knocked out via a standard biolistic bombardment, homologous recombination, and selection protocol (57). In brief: PCR was used to amplify the 5’ and 3’ flanking regions (1.5–2 kb each) for each gene. These amplicons were subsequently subcloned into the *Sac*I and *Xho*I sites that flank a neomycin resistance cassette (*Neo4*), thus granting the cassette homology arms to replace the endogenous gene. These vectors were linearized by *Kpn*I and *Sap*I and transformed into CU428 cells by biolistic transformation. Biolistic transformations were as described previously (58), with the following modifications: gold particles were prepared as recommended with 15 μg of total linearized plasmid DNA. To select positive transformants, paromomycin was added 4 hours after bombardment to cultures that had been shaking at 30°C. Transformants were selected in 120 ug/mL paromomycin, and CdCl2 was added at 1 μg/mL to induce *Neo4* expression. Putative transformants were identified after 3 days of selection. These were then serially transferred daily in increasing amounts of paromomycin for at least 4 weeks before further testing.

### Dibucaine mucocyst secretion assay to experimentally validate new mucocyst gene knockouts

*T. thermophila* cells (wildtype or knockout) were grown to a density between 4 × 10^5^ and 6 × 10^5^ cells/mL and washed once with 10 mM Na-HEPES (pH 7.2) after being pelleted for 30 seconds at 400 × *g* in a clinical centrifuge. Loose cell pellets (concentrated ∼10-fold relative to the initial culture) were stimulated for 30 s by addition of 2.5 mM dibucaine (final concentration). The cells were then diluted at least five-fold with 10 mM Na-HEPES (pH 7.2) and centrifuged at 1, 200 × *g* for 1 min in 15 mL conical tubes. After the centrifugation, we imaged the two-layer pellet, with cells overlaid by flocculent extruded mucocyst contents, to determine the strains’ relative capability to secrete mucocysts.

## Results

### Uniting insights from microarray and RNA-seq datasets

In order to draw comparisons between gene expression patterns in the disparate microarray and RNA-seq datasets, we processed them into a *lingua franca* of normalized gene expression. The microarray expression dataset covered bulk growth, starvation, and conjugation conditions (12, 13, 15), and the RNA-seq expression dataset covered 1.5 cell cycle–synchronized mitotic cycles (26). We filtered both datasets, removing batch effects, noisy expression, and unreplicated samples (Supplementary Figures 1 and 2). After the quality control steps, 20, 428 genes remained in the microarray dataset and 23, 113 genes remained in the RNA-seq dataset (the total gene number in *T. thermophila* is 27, 494) (27) (Table 1). After normalization, we scanned over 5 different distance metrics (Euclidean, Manhattan, context likelihood of relatedness (CLR), angular, and linear correlation (13, 32, 59)), nearest neighbors ranging from 2-12, and scanning the Leiden clustering resolution parameter between zero and one (Supplementary Figures 3-7) (24). Using Pareto optimization (30), we settled on Manhattan distance, three nearest neighbors, and resolution parameter (r = 0.005) as the most effective (Supplementary Figure 3). The Pareto optimization checked for modularity (60), fraction of clusters with significantly enriched functional terms, and cluster size interquartile range (Supplementary Figures 3-7). To determine the functional term enrichment, we first used eggNOG and InterProScan to annotate all the genes in our dataset based on orthologous groups and protein domains (21, 22, 28) (Table 1). In each cluster, the enrichment of each functional term was calculated against its background abundance in the genome using a modified Fisher’s Exact Test and Bonferroni correction against multiple hypothesis testing (45, 46) (Table 2).

**Table 1.** Annotation statistics of the microarray and RNA-seq datasets.

|  | Microarray | RNA-seq |
| --- | --- | --- |
| Total Number of Genes | 20428 | 23113 |
| Fraction Genes with COG Category Terms | 0.55 | 0.52 |
| Fraction Genes with GO Terms | 0.08 | 0.07 |
| Fraction Genes with KEGG KO Terms | 0.32 | 0.29 |
| Fraction Genes with EC Terms | 0.14 | 0.13 |
| Fraction Genes with PFAM Terms | 0.49 | 0.45 |
| Fraction Genes with InterPro Terms | 0.65 | 0.63 |

**Table 2.**
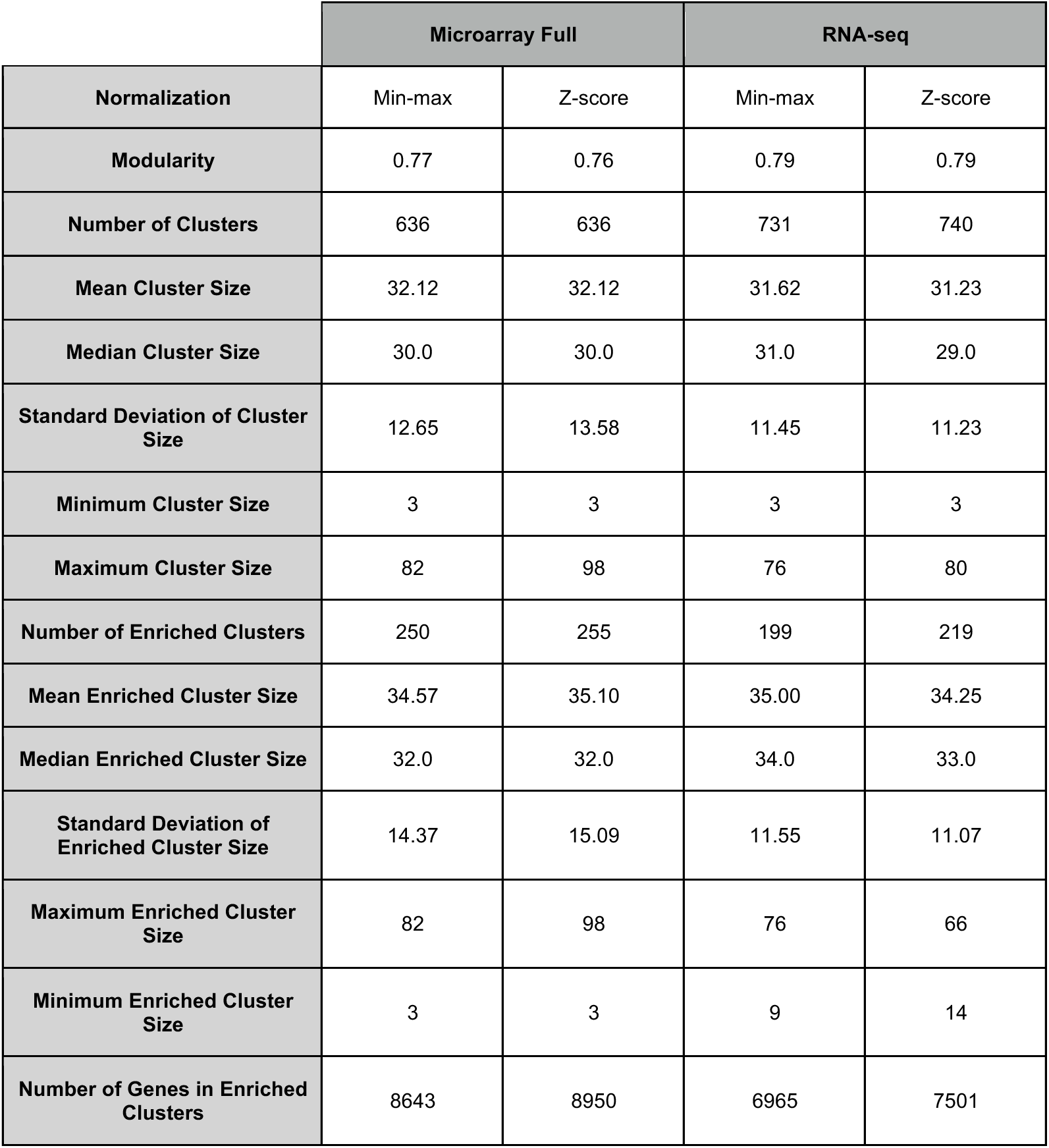
Normalization specific cluster and enrichment statistics of the optimal microarray and RNA-seq partitions.

After all these steps, we simulated the null hypothesis that each gene expression profile was completely unrelated to other gene expression profiles with two methods: (1) expression value scrambling within each gene, or (2) generating simulated expression profiles each gene that supported an evenly distributed hypercube of values (Figure 1). For each method, we ran 1000 independent simulations and found that the normally distributed modularity values corresponding to the null hypothesis were statistically significantly lower than the partition modularity for the chosen optimal partitions (p < 0.005 by two-tailed t-test), which indicated that our parametrization identified significant gene co-expression modules (Figure 1A-D). For both the microarray and RNA-seq datasets, and for both normalization strategies, we hierarchically clustered the co-expression modules around their centroids, allowing us to plot gene co-expression modules by their relative similarity (Figure 1E-H). These heatmaps reveal a whole-transcriptome view of gene expression across all the assayed conditions, and each condition has a corresponding co-expression module that reaches either its minimum or maximum at that point.

**Figure 1.**
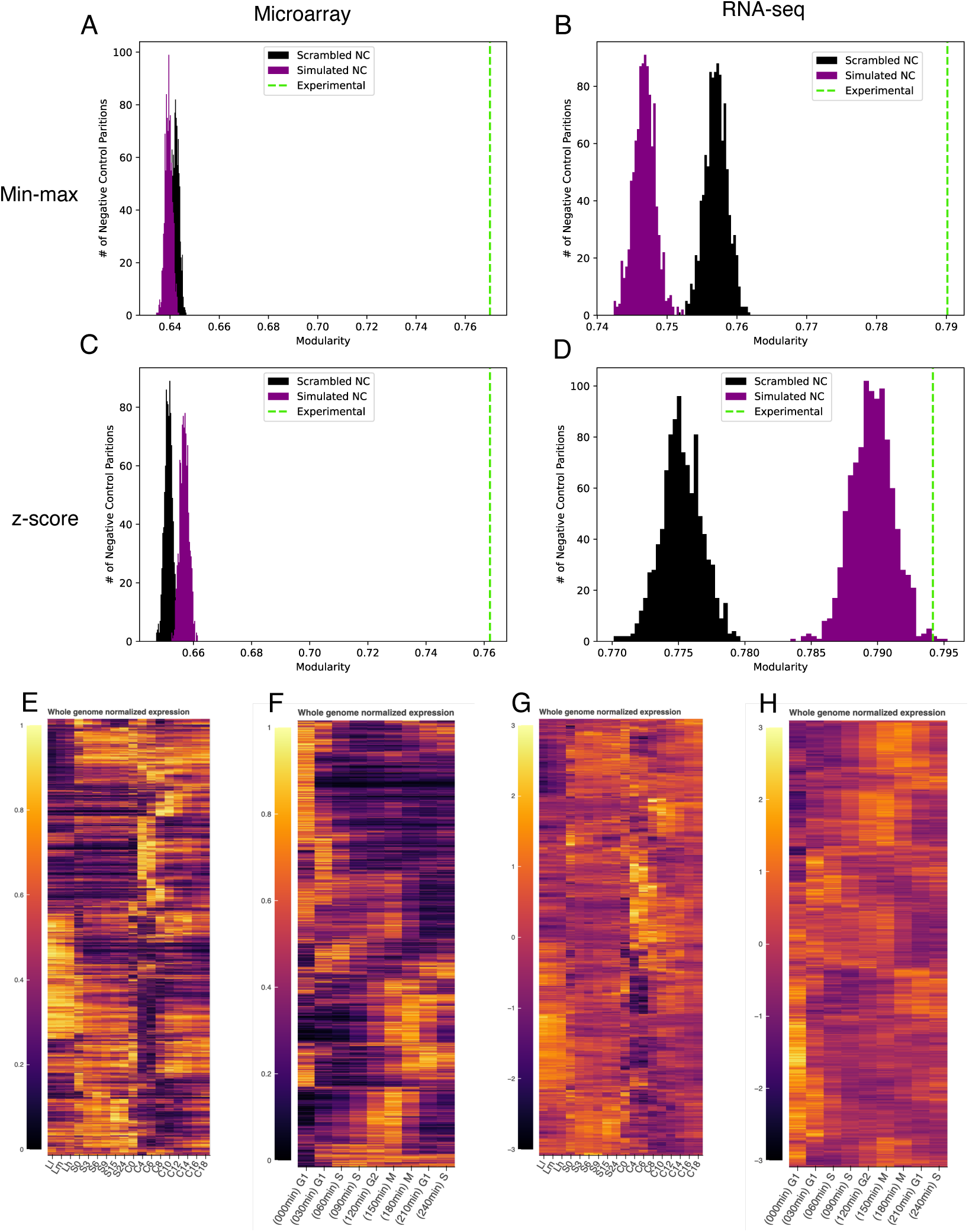
Optimal parameterization significance testing for each dataset/normalization scheme and illustrations of optimal experimental partitions. Histograms illustrating modularity distributions for computational negative control (NC) partitions compared to the experimental partition created from the optimal parameterization. The computational negative controls based on scrambled data are in black, the computational negative controls based on a simulated hypercube with uniform data distribution are in purple, and the modularity value for the optimized partitions are indicated by the dashed green line. (A) The computational negative control comparison for the min-max normalized microarray dataset. (B) The computational negative control comparison for the min-max normalized RNA-seq dataset. (C) The computational negative control comparison for the z-score normalized microarray dataset. (D) The computational negative control comparison for the z-score normalized RNA-seq dataset. In each case, the modularity for the optimized clustering of the real data was statistically significantly greater than in either negative control (p < 0.005). Heatmaps illustrating the optimal partitions generated from (E) the min-max normalized microarray dataset, (F) the min-max normalized RNA-seq, (G), the z-score normalized microarray, and (H) the z-score normalized RNA-seq datasets. Modules of gene expression profiles are ordered by hierarchical clustering of the module centroids using average linkage. Each row of a given heat map corresponds to one gene’s expression. In (E) and (G), the x-axis denotes the different phases of the *T. thermophila* life cycle: low density logarithmic growth (Ll), medium density logarithmic growth (Lm), high density logarithmic growth (Lh), 0-24 hours of starvation (S0-S24), and 0-18 hours of conjugation (C0-C18) (12). In (F) and (H), the x-axis denotes the stages of the mitotic cell cycle and corresponding timepoints for sampling.

### Mucocyst biogenesis genes are recovered in untargeted bioinformatic analysis

With the gene co-expression being normalized and computed in the same way for the two datasets, we were able to test the degree of their agreement. We focused on mucocyst biogenesis, a cellular process that we have previously studied, including with the use of inferences from gene co-expression (16, 18, 20). For our current analysis, we determined which clusters in both the microarray and RNA-seq datasets were enriched for 33 genes that have been previously experimentally verified to be involved in mucocyst biogenesis or secretion (Supplementary File 1). Using the min-max normalized data, we found clusters that are enriched for these experimentally verified genes: six in the microarray co-expression dataset (m002, m003, m004, m005, m006, m378, totaling 182 genes, Figure 2A) and four in the RNA-seq co-expression dataset (m040, m194, m219, and m294, totaling 104 genes, Figure 2B). Using the z-score normalized data, we found clusters four such clusters in the microarray co-expression dataset (m169, m171, m172, m424, totaling 172 genes, Supplementary Figure 9A) and four in the RNA-seq co-expression dataset (m632, m634, m636, m679, totaling 144 genes, Supplementary Figure 9B).

**Figure 2.**
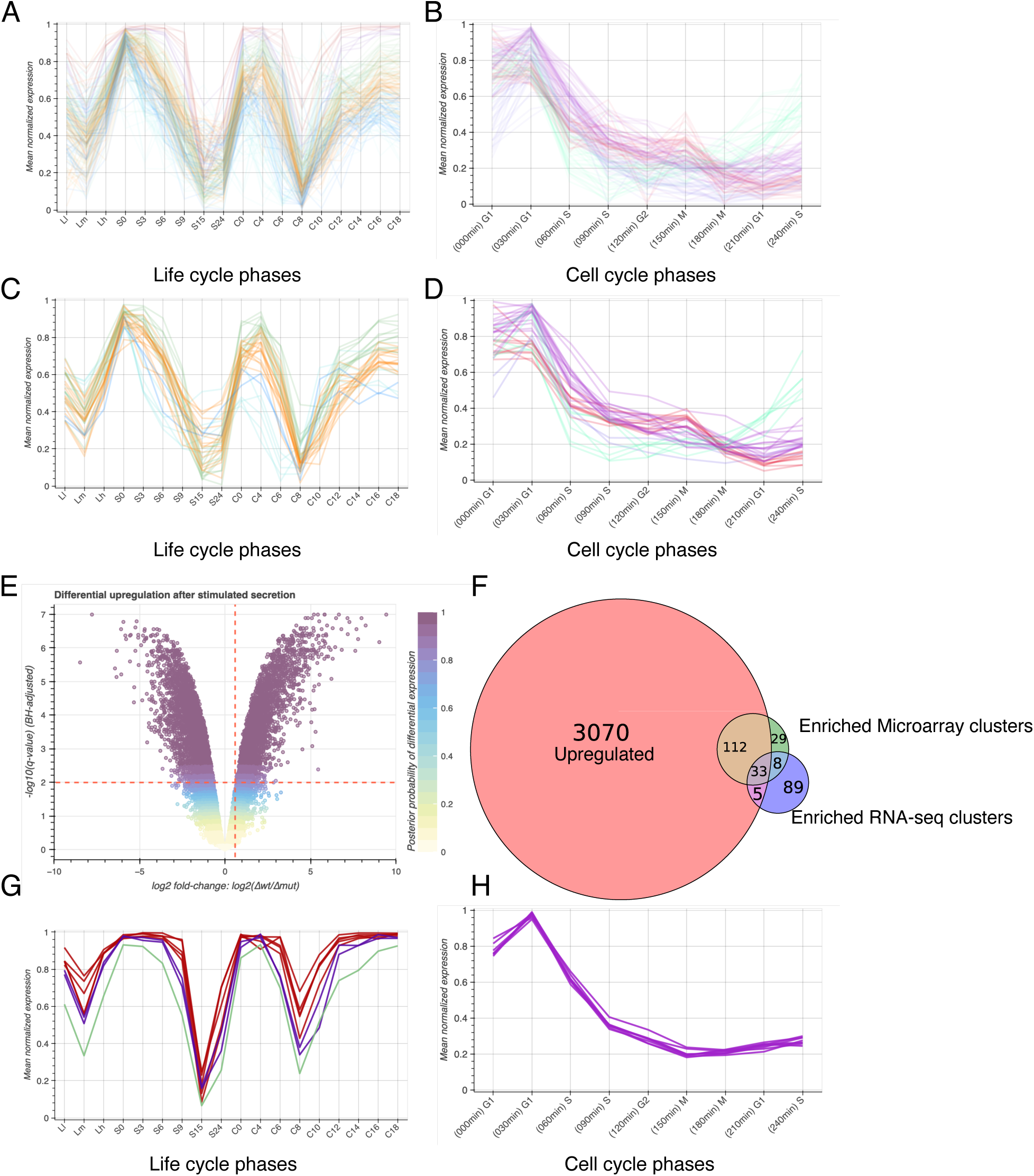
Enrichment, differential expression, and overlap of experimentally validated mucocyst-associated and differentially expressed, upregulated genes. Min-max normalized expression profiles for genes in (A) the six microarray and (B) the four RNA-seq clusters significantly enriched for experimentally validated mucocyst-associated genes as well as the 33 genes overlapping between the upregulated, enriched microarray clusters, and enriched RNA-seq clusters in (C) the microarray and (D) the RNA-seq datasets. (E) Volcano plot illustrating differential expression of each gene represented in the microarray dataset over one hour in the MN173 mutant relative to the wild type *T. thermophila*. Thresholds are represented by blue dashed lines (q < 0.01 and fold-change > 1.5). All genes that passed the thresholds have a Bayesian posterior probability of differential expression greater than 80%. (F) Venn diagram describing the overlapping genes in the enriched microarray clusters, enriched RNA-seq clusters, and the set of upregulated genes with min-max normalization. Min-max normalized expression profiles for genes that are co-expressed in the microarray and RNA-seq datasets, but not detected in the upregulated dataset: (G) gene expression in the microarray profiles and (H) gene expression in the RNA-seq profiles.

To determine whether this agreement between the two datasets has biological significance, we experimentally assessed which genes are upregulated after we stimulated massive mucocyst secretion, when the cells are induced to synthesize a large cohort of these organelles. For this analysis, we compared a wildtype strain (CU428) to a mutant (MN173) that produces mucocysts but is incapable of releasing them, and which therefore does not induce new mucocyst synthesis upon stimulation (61). This was a microarray assay that we processed in the same way as described above (Supplementary Figure 8). Focusing on min-max normalized data, of the 3220 genes that were differentially upregulated in this experiment (Figure 2E), 112 are shared with the co-expressed clusters in the microarray dataset alone and five are shared with the ones in the RNA-seq dataset alone (Figure 2F-H). 33 genes are shared across the differential upregulation and the two co-expression datasets (Figure 2F).

In Figure 2F, the 33 gene intersection of the three datasets includes 13 experimentally verified genes (Supplementary File 1, “min-max triple agreement” tab), which is a statistically significant enrichment of experimentally verified genes relative to the background of all genes in Figure 2F (p < 1 x 10^-6^ by a two-tailed Fisher’s Exact test). The five gene intersection of the RNA-seq co-expression and differential upregulation alone contains no experimentally verified genes (Supplementary File 1, “min-max upreg & RNA-seq” tab; p = 1). The 112 gene intersection of the microarray co-expression and differential upregulation alone contains 8 experimentally verified genes (Supplementary File 1, “mix-max upreg & microarray” tab; p < 1 x 10^-6^). The eight gene intersection of the microarray and RNA-seq co-expression alone contains seven experimentally verified genes (Supplementary File 1, “min-max microarray & RNA-seq” tab; p < 1 x 10^-6^). Thus, 28 of the 33 genes (84%) that were experimentally verified to be involved in mucocyst biogenesis prior to this analysis were recovered, all of which are found in the microarray co-expression dataset and shared agreed upon by at least one of the two other datasets. We obtained analogous results when starting with z-score normalized data (Supplementary Figure 9 and Supplementary File 1, “z-score” tabs).

Figure 2G-H shows the expression profiles of the 8 genes at the intersection of the microarray and RNA-seq co-expression datasets. This list includes GRL1, 3, 4, 5, 7, and 8 and GRT1, which are all known to be mucocyst cargo proteins (58, 62–64). The other gene is TTHERM_00537380, which is unnamed, unannotated, and lacks any orthologs or protein domains that were identified by eggNOG or InterProScan (Supplementary File 1).

For 10 of the genes that are co-expressed with the 33 previously verified genes, we performed new knockout experiments confirming their role in mucocyst biogenesis (Figure 3). Six of these genes have been previously implicated to be part of the Mucocyst Docking and Discharge complex by co-immunoprecipitation but have not been genetically assayed (Figure 3A) (5). The remaining four had not been previously studied, but we selected them based on their co-expression clusters and for their putative annotations as proton-pumping pyrophosphotases, which have been implicated in mucocyst and trichocyst biogenesis (Figure 3B) (65–67). In our knockout experiments, the loss of each of these ten genes resulted in a mucocyst secretion defect, as evidenced by the loss of the mucosal layer over the cell pellets after dibucaine treatment.

**Figure 3.**
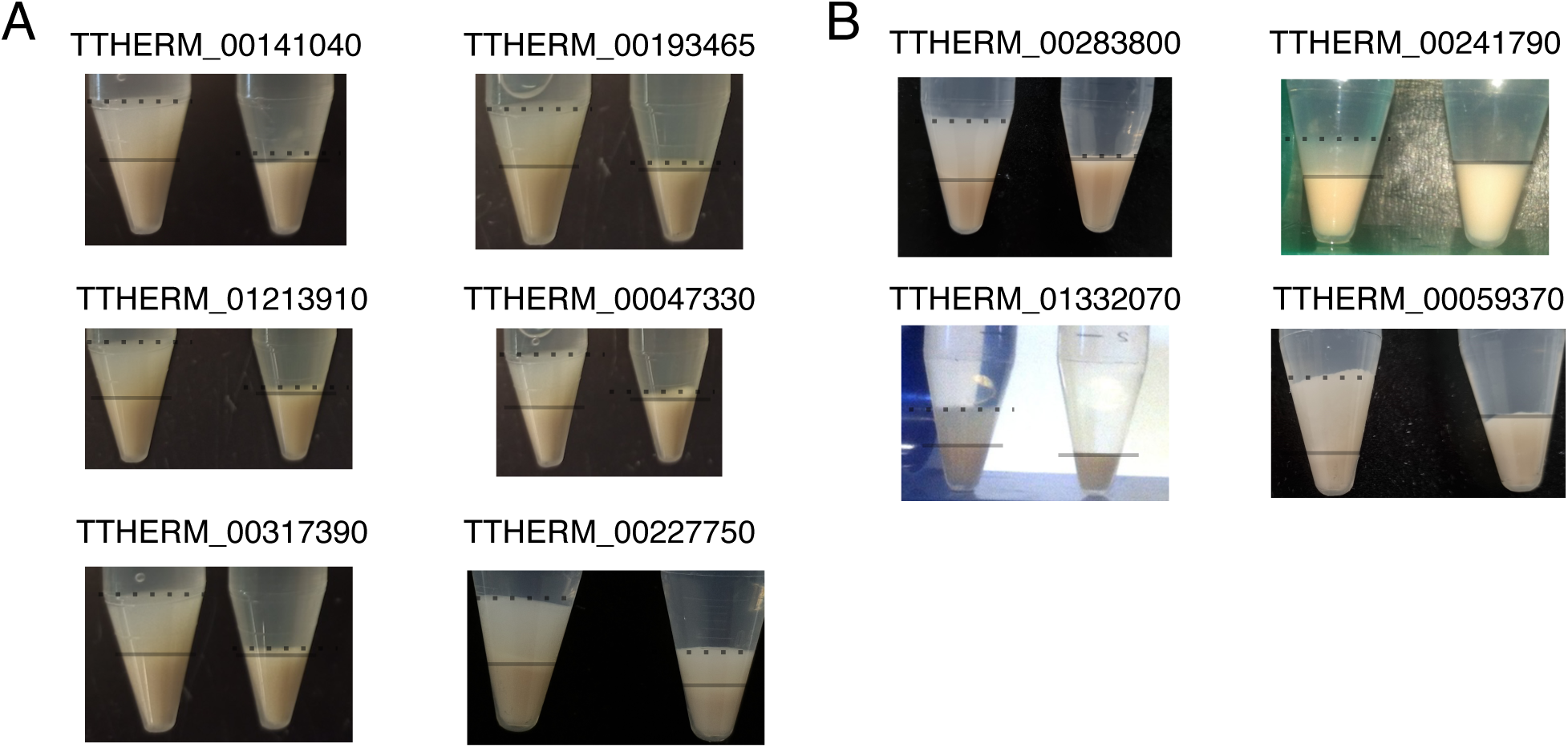
Experimental validation of ten genes that are suggested to be mucocyst-associated by our co-expression analysis. (A) Genes that co-immunoprecipitated as members of the Mucocyst Docking and Discharge protein complex (TTHERM_00141040, TTHERM_00193465, TTHERM_01213910, TTHERM_00047330, TTHERM_00317390, and TTHERM_00227750) (5). (B) Four genes that were knocked out solely on the basis of our co-expression inference (TTHERM_00283800, TTHERM_00241790, TTHERM_01332070, and TTHERM_00059370). For each gene, the left tube shows the wildtype response to dibucaine as evidenced by a flocculent layer of mucus overlying the cell pellet after centrifugation. The boundary of the cell pellet is denoted by the solid line, and the boundary of the mucus layer is denoted by the dotted line. The right tube in each panel displays the phenotype of strains with the respective genes genetically knocked out. Each has a defect in mucocyst release in response to the dibucaine treatment.

*The topology of the T. thermophila gene network reveals other functionally enriched modules* The previous two studies of the *T. thermophila* gene co-expression landscape identified relatively large modules with significantly enriched functional terms: the RNA-seq study identified 3032 genes as cell cycle-regulated that were divided into 10 clusters, only four of which had significantly enriched functional terms (26). The microarray study reported 55 co-expression clusters for the full genome, but did not report a statistical analysis of functional enrichment (13). However, both studies found functionally associated genes clustering together––most prominently histone-, proteasome-, and ribosome-associated genes. We set out to assess whether our new analysis reproduces or expands on these findings (Figure 4).

**Figure 4.**
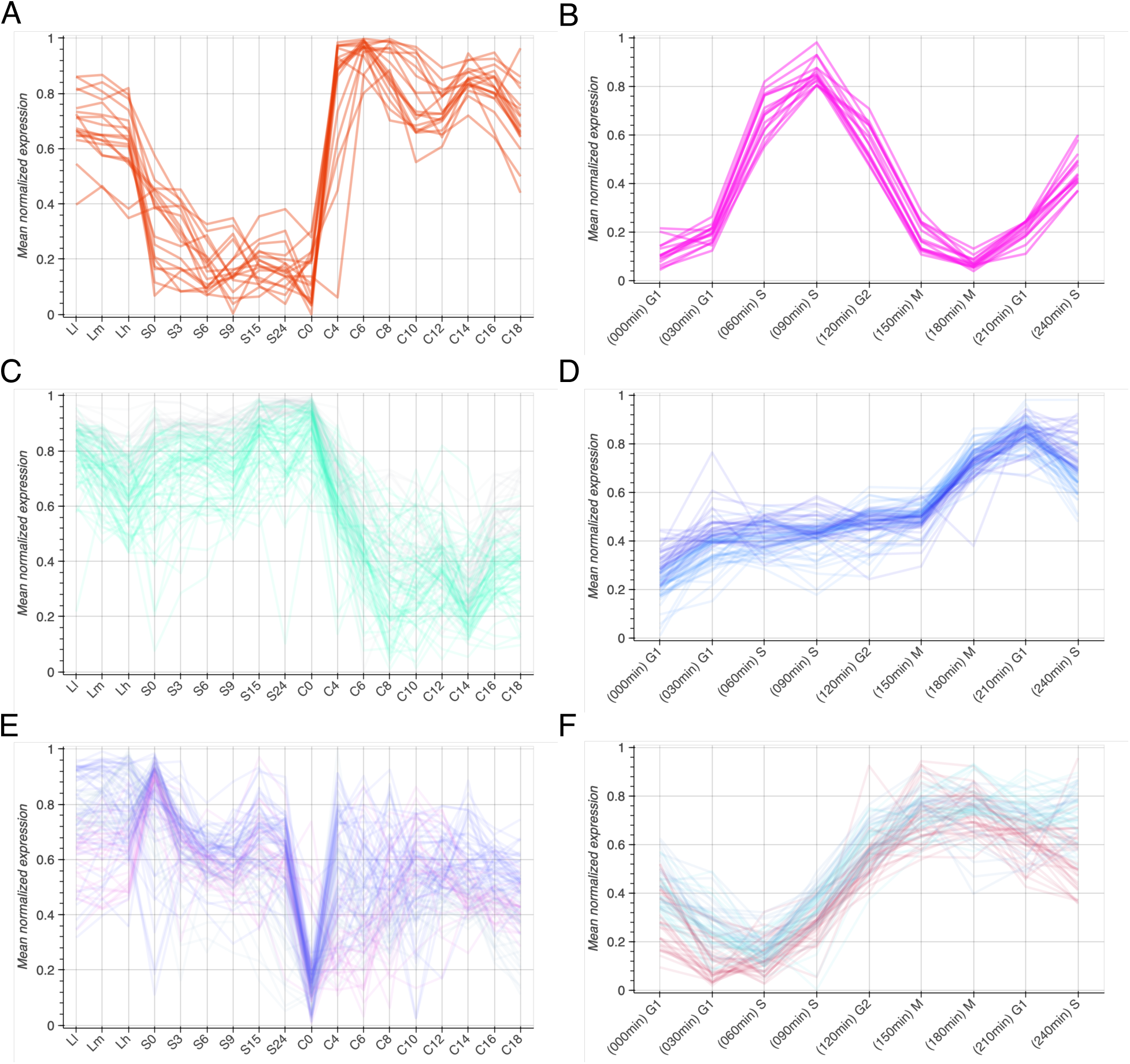
Min-max normalized expression profiles of clusters significantly enriched for (A-B) histone, (C-D) ribosome, and (E-F) proteasome functional annotation terms in the microarray (left column) and RNA-seq (right column) analyses. In each case, the same number of clusters come up in the two datasets: one for histone-associated profiles, two for ribosome-associated profiles, and three for proteasome-associated profiles. The histone-associated profiles are characterized by low expression during starvation and high expression during growth and conjugation (A) and high expression during the S-phase of the cell cycle (B). Ribosome-associated profiles are characterized by high expression during growth or starvation and low expression during conjugation (C). In the RNA-seq expression dataset, the ribosome-associated profiles appear to be at a minimum during the first G1 phase and at a maximum at the second G1 phase, indicating that in this experiment they are not following the cyclicity of the mitotic cell cycle (D). The main characteristics of the proteasome-associated co-expression pattern are a sharp loss of expression at the beginning of conjugation (E) and a peak of expression during mitotic division (F).

In the case of histone-associated genes, the min-max microarray co-expression analysis identified one cluster of 18 genes (module m179), and the min-max RNA-seq co-expression analysis identified one cluster of 15 genes (module m721) (Figure 4A-B). The intersection of these two gene sets comprises six genes, five of which were previously annotated as histone components: TTHERM_00790792 (HTA1), TTHERM_00633360 (HTB1), TTHERM_00498190 (HHF1), TTHERM_00316500 (HTA2), TTHERM_00283180 (HTB2), and TTHERM_00143660 (HTA3) (Supplementary File 2, “AB overlap”). Additionally, the microarray co-expression cluster identifies five members of the MCM helicase and a chromatin-associated protein: TTHERM_00554270 (MCM2), TTHERM_00092850 (MCM3), TTHERM_00277550 (MCM4), and TTHERM_00448570 (MCM6), TTHERM_00011750 (putative MCM7), and TTHERM_00729230 (IBD1) (Supplementary File 2, “Fig4A”). The RNA-seq co-expression cluster also includes more histone- and chromatin-associated genes, such as: TTHERM_00823720 (HHO1), TTHERM_00660180 (HMG1), TTHERM_00257230 (HMGB2), TTHERM_00189170 (HHF2), and TTHERM_00571055 (HHT1) (Supplementary File 2 “Fig4B”).

For the ribosome-associated genes, the microarray co-expression analysis identified two clusters with functional enrichment, modules m601 and m602, comprising 105 genes (Figure 4C). The RNA-seq analysis also identified two clusters, modules m458 and m460, comprising 82 genes (Figure 4D). The overlap between the two consists of 49 genes, each one of which is annotated as a ribosomal gene (Supplementary File 2, “CD overlap”). Similarly, the proteasome-associated genes separate into three co-expression profiles in both the microarray analysis (modules m374, m375, and m453; 102 genes total) and the RNA-seq analysis (m467, m470, m473; 87 genes total) (Figure 4E-F). The intersection between these gene sets contains 22 genes, 21 of which are annotated as proteasomal components (Supplementary File 2, “EF overlap”). The exception is TTHERM_00600110 (TTN1), which is a nuclease (68). Curiously, the ribosomal co-expression profiles do not follow the periodicity of the mitotic cell cycle (Figure 4D), unlike the histone (Figure 4B) and proteasome (Figure 4F) co-expression profiles.

### The Interactive Tetrahymena Gene Network Explorer (TGNE)

Given that our analysis appears to be broadly informative for *T. thermophila* cell biology, we developed an interactive tool for reproducing our investigations for any gene or pathway of interest (Figure 5). The TGNE is a standalone HTML file that contains all the data, making it portable and requiring no maintenance. The microarray version is 292.1 MB, and the RNA-seq version is 106.2 MB. The TGNE works in any web browser that supports webGL 2.0 (e.g., Chrome 56+, Firefox 51+, Safari 15+, and Opera 43+), and every plot in the tool is interactive. The annotation table with the selected genes, as well as the functional term enrichment data for the corresponding modules, can be downloaded using the buttons at the upper right corner (Figure 5H). The functional term enrichment data for each module in each variant of the TGNE is available in Supplementary File 3. The interactive dashboards for the microarray and RNA-seq variants of the TGNE are available as Supplementary File 4 (microarray version) and Supplementary File 5 (RNA-seq version).

**Figure 5.**
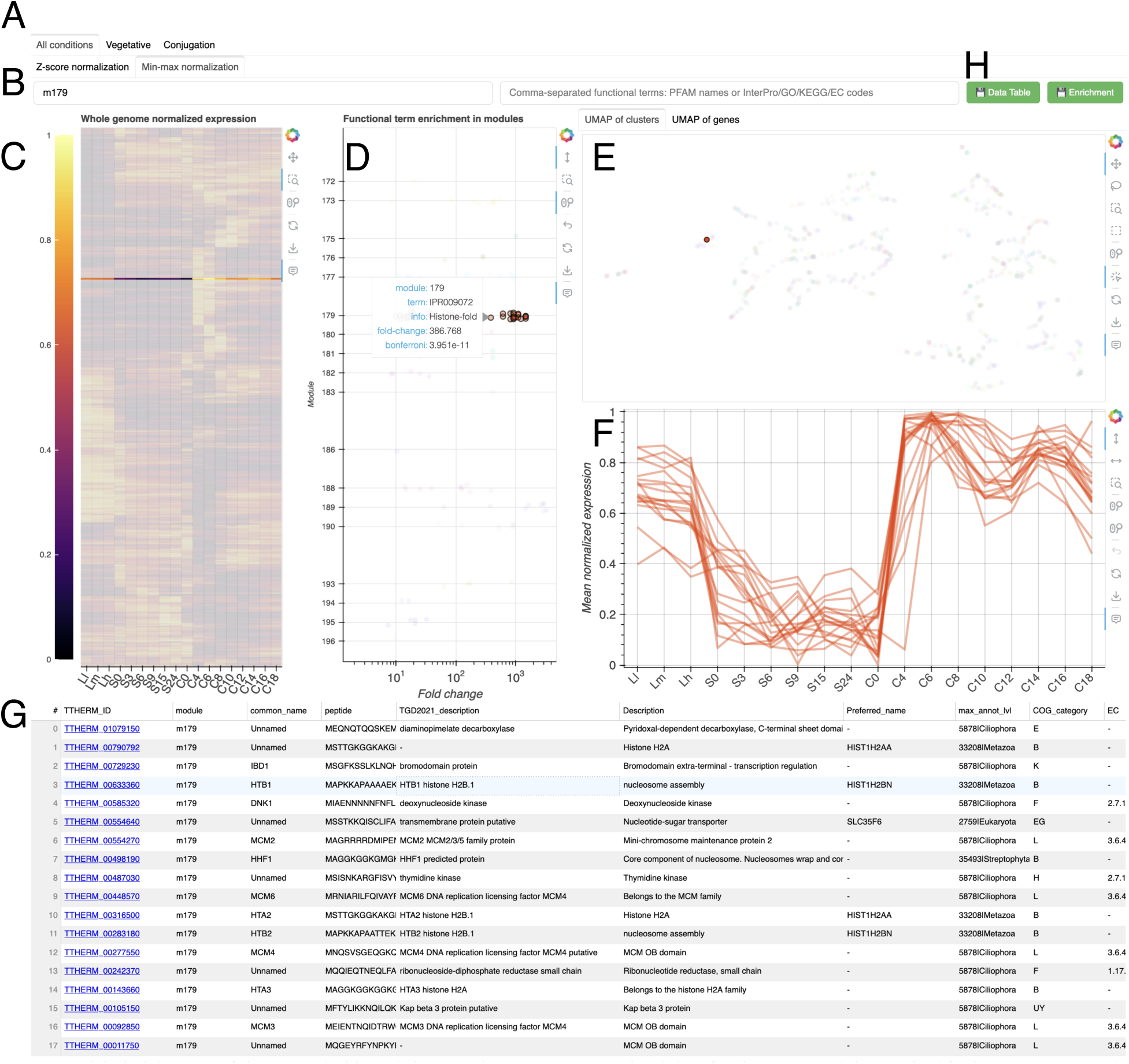
A labeled diagram of the TGNE dashboard showing the min-max normalized data for the gene module enriched for histone-associated genes. (A) The “Conditions Selection Tabs” are exclusive to the microarray dashboard and allow the user to specify which life cycle phases are included within the input data to the clustering pipeline: the entire profile, just the vegetative conditions, or just the conjugative conditions. The “Normalization Selection Tabs” allow the user to select which normalization technique should be used on the input data: z-score or min-max. (B) The search bars are text fields that can be used to select genes based on their annotations. The left search bar allows searches for TTHERM_ID, common names, descriptions, and module number. The right search bar allows searches for functional annotation terms or codes, specifically: PFAM names or GO/KEGG/InterPro/E.C. alphanumeric codes. Here, “m179” was used as the search term to select the entire module that is enriched for histone-associated functional terms. (C) The heatmap representation of the normalized expression of all genes across all conditions, as in Figure 2E. The selected module is highlighted, and the unselected genes are grayed out. (D) This plot shows all modules with significantly enriched functional terms, which are the same terms as those that can be searched using the right-hand search bar. As with the heatmap, when a certain module is selected, the others are grayed out. Moving the cursor over any of the circles in the plot displays the enriched term, its fold-change relative to the genome background, and the Bonferroni-corrected p-value. Here, the indicated circle represents the InterPro term “IPR009072”, which corresponds to “Histone-fold”. This term is 386 times over-represented in this cluster relative to the genome background, with a Bonferroni-corrected p-value of ∼4 x 10^-11^. (E) An interactive UMAP representation of the gene expression with one tab showing the UMAP embedding of each cluster and the other tab showing the UMAP embedding of each gene. Selected genes and modules are highlighted, while unselected ones are grayed out. Clicking on any circle or selecting them with one of the tools to the right of the plot selects those module(s) or gene(s) for display. (F) The graph for displaying the expression profiles of the selected genes. This is an equivalent representation of the data in the heatmap. (G) The annotation table. When genes are selected, their annotation information based on the published *T. thermophila* genome, eggNOG, and InterProScan is populated into this table. Columns after the EC terms are not displayed in this figure. (H) The download buttons. The annotation table and functional enrichment information for the selected genes/modules can be downloaded as tab-separated files using these two buttons.

## Discussion

Our results show that, at least in the four cases we explored, there is significant agreement between co-expression patterns in two disparate experiments: one measuring gene expression using microarrays across growth, starvation, and conjugation and one measuring gene expression using RNA-seq in cell cycle-synchronized cells, which were generated fifteen years apart in different laboratories (12, 26). The fact that we were able to bring the data into a shared framework that bridges these experimental and temporal differences is an indication that old data do not need to lie fallow. Crucially, our approach relies on careful quality filtering, normalization, systematic parameter optimization, and computational negative controls. The basis for our reanalysis was to align the probes/reads against the newest model of the genome and then normalize the expression data such that the resulting co-expression clusters would be translatable between datasets. Each dataset was filtered to remove batch effects, unreplicated gene expression, and noise (Supplementary Figures 1 and 2). We normalized each dataset in two ways to satisfy two different perspectives on the data. Min-max normalization linearly scales the data between zero and one, emphasizing only the shape of the given expression profile. In contrast, z-score normalization incorporates both the shape and the magnitude of expression within each profile. We hope that our work can be a roadmap for using co-expression data to identify functionally associated genes in other systems.

Our comparison of metrics for difference (or similarity) between gene expression profiles showed that the Manhattan distance (also termed the L1 Norm) performs as well or better than the other metrics, which is in line with prior literature on overcoming the “curse of dimensionality” (Supplementary Figures 3-7) (69, 70). For all distance metrics except Context Likelihood of Relatedness (CLR), three nearest neighbors was the best parametrization for Leiden clustering (Supplementary Figures 3-7) (24). Notably, the CLR distance metric worked significantly differently for the microarray and RNA-seq expression datasets, unlike any of the other metrics (Supplementary Figure 5A-B), which indicates that it would be inappropriate for bridging the two datasets. The previous *Tetrahymena* gene network landscape employed CLR to detect co-expression clusters, so our analysis is fundamentally distinct (13). The other four distance metrics appear to be largely equivalent in terms of the resulting modularity, fraction of clusters with enriched functional terms, and interquartile range for cluster size (Supplementary Figures 3, 4, 6, and 7). We chose the Manhattan distance for our subsequent analyses because it gave the most similar clustering statistics between the microarray and RNA-seq datasets, but it is possible that the other distance metrics could reveal subtle differences in the detected co-expression patterns.

The Leiden algorithm generates flat clusters, as opposed to hierarchical ones. Consequently, we do not report the relative pairwise relatedness of genes, unlike the previous *Tetrahymena* gene network analysis (13). This approach allowed us to calculate statistically significantly enriched terms for ∼40% of clusters (Table 2). However, the interactive TGNE dashboard can be used to glean inter-cluster relatedness based on the heatmap, which is hierarchically sorted based on each cluster’s centroid in geometrical space, as well as the UMAP embeddings and shared annotation enrichment terms (Figure 5C-E).

We specifically chose to incorporate modularity and the fraction of enriched clusters in the three-dimensional Pareto optimization to optimize for both mathematical and biological significance, respectively. A higher relative modularity indicates that there are more intra-cluster paths (i.e., more relatedness between gene expression profiles in the same cluster) and fewer inter-cluster paths (i.e., less relatedness between genes in different clusters) (60). The fraction of enriched clusters corresponds to the proportion of clusters that have statistically significantly enriched functional annotations relative to the distribution of annotation terms in the genome. Modularity and functional enrichment fraction are not sufficient to avoid partitions where there are either several large clusters containing all genes or where the majority of clusters are tiny, neither of which would be biologically informative. Optimizing the interquartile range for cluster size allowed us to avoid parameter settings that resulted in either of these situations. From the Pareto optimal set of partitions, the first optimal partition with an interquartile range of cluster sizes greater than ten was chosen, as this was the point where cluster sizes were still small enough to be manually verified.

Our approach enabled us to perform computational negative controls to assess whether our clustering performs better than the null hypothesis––that there is no co-expression network in the data (Figure 1). This methodology allowed us to analyze all the datasets in the same way and ensure the validity of our chosen partitions. Interestingly, the two variants of the negative control distributions for each dataset were never completely superimposed. The min-max normalized datasets illustrated scrambled negative control distributions which had a higher average modularity than the simulated distributions. The opposite was true for the z-score normalized datasets. While the use of computational negative controls in co-expression studies has previously been employed qualitatively or propounded on theoretical grounds (1, 29–31), our treatment of these *T. thermophila* datasets is the most systematic that we are aware of. Furthermore, the consistent modularity scores of our Pareto-optimized partitions and the performance of the negative controls gave us confidence to draw comparisons between the co-expression patterns in the microarray and RNA-seq datasets.

Our primary goal for the TGNE was to develop a tool for generating testable hypotheses about *T. thermophila* cell biology. To evaluate its effectiveness, we used it to revisit the biogenesis of an organelle in *Tetrahymena* that we have previously studied: the mucocyst (Figure 2). Using the TGNE, we found clusters which were enriched for the 33 genes that are experimentally known to be associated with mucocysts (i.e., genes that either are required for mucocyst biogenesis/secretion, localize to mucocysts, or both) in both the RNA-seq and microarray datasets (Figure 2A-B). We compared these genes against our differential upregulation experiment, which assessed gene expression in wildtype or secretion-null mutant cultures after stimulating mucocyst release. The degree of agreement between co-expression and differential upregulation patterns, as displayed in the intersections of the Venn diagrams in Figure 2F and Supplementary Figure 9F, gave us confidence that our co-expression clusters contain novel genes that are important for mucocyst biogenesis. We generated new knockouts for 10 genes (Figure 3): six were previously co-immunoprecipitated with the Mucocyst Docking and Discharge complex (Figure 3A) (5); four were unstudied but had putative annotations as proton-pumping pyrophosphotases, which have been implicated in ciliate membrane trafficking (Figure 3B) (65–67). Each knockout had a mucocyst secretion defect, and after this initial confirmation, these genes will be the subject of detailed future studies.

Importantly, each intersection in the Venn diagram in Figure 2F and Supplementary Figure 9F indicates other clear candidates for genes involved in mucocyst biogenesis that have not been previously studied (Supplementary File 1). These intersections include four orthologs to the *Paramecium tetraurelia* trichocyst cargo proteins (TTHERM_00321725, TTHERM_00773710, TTHERM_00697290, and TTHERM_00773700) and eight members of an expanded gene family that shares a beta/gamma crystallin domain with known mucocyst cargo genes (TTHERM_00585170, TTHERM_00471040, TTHERM_00038880, TTHERM_00558350, TTHERM_00570550, TTHERM_01002860, TTHERM_01002870, and TTHERM_00989430) (71–73). There are also two genes that are known to be essential for trichocyst secretion: ND6 (TTHERM_00410160) and ND9 (TTHERM_00938850) (74, 75).

Furthermore, eight genes which were co-expressed in both the microarray and RNA-seq datasets but were not induced upon regranulation (Figure 2F) include GRL1 (TTHERM_00527180), GRL3 (TTHERM_00624730), GRL4 (TTHERM_00624720), GRL5 (TTHERM_00378890), GRL7 (TTHERM_00522600), GRL8 (TTHERM_01055600), GRT1 (TTHERM_00221120), and TTHERM_00537380 (which has no annotation or clear orthologs outside the ciliates available) (Supplementary File 1). Of these, the GRLs and GRT1 are known mucocyst cargo genes (51, 58, 62, 63, 71, 72). Every gene in this set has an average log2 intensity >15.5 on the microarray, apart from the unnamed gene, which is >14.9. Given the 16-bit detection camera of the system that was used for the microarray-based experiments, there was not enough dynamic range to detect further upregulation of these genes (Supplementary File 6). However, we have previously detected upregulation of GRL1, GRL3, and GRL4 during regranulation by qPCR and Northern blot in the same experimental framework (76). Thus, we expect that if the differential upregulation experiment were performed using RNA-seq instead of microarrays, the majority of these genes would also be in the triple-intersection of the Venn diagram. Of note, the completely unannotated TTHERM_00537380 was over-represented in the constitutive secretome of the *T. thermophila* SB281 mutant, which is also the case for mucocyst cargo proteins (Supplementary File 1) (51, 77, 78). The fact that it shares a strong co-expression profile with verified cargo genes and is in this secretome makes TTHERM_00537380 a prime candidate for future study.

A consequence of our clustering approach is that it highlights statistically significant, but qualitatively subtle, differences in expression patterns among well-characterized complexes or functionally related genes. One intriguing example of this phenomenon is found within the nine-member gene family called GRL, for <u>Gr</u>anule <u>L</u>attice. All GRL products are structurally related secretory proteins that are co-packaged within mucocysts, and early analyses of transcriptomic data revealed that the GRL genes are highly co-regulated. However, while most GRL proteins are likely to be required to form the physical core of the mucocyst, *GRL6* appears to play a distinct regulatory role, as yet poorly understood (62). Remarkably, in our current analysis *GRL6* partitions into a different cluster from the other GRL genes, potentially reflecting this functional divergence (Supplementary Figure 10). We posit that, even in the absence of any other data, the separate clustering of closely related genes may provide hints of functional diversification. More broadly, in cases where specialized cell biological structures or pathways in *T. thermophila* were co-inherited by other ciliates or the relatively closely related dinoflagellates and apicomplexans, the transcriptional clusters detected in the TGNE may help to uncover novel features within this deep lineage.

The efficacy of bioinformatic approaches like ours is necessarily limited by the input datasets. Even though we were able to “modernize” the microarray data by aligning it to the newest genome model and applying more quality control, microarray datasets are inherently limited by both maximum and minimum signal intensities. This results in a plateau effect in gene expression profiles in which high expression levels reach a signal ceiling and low expression levels fall below detection thresholds. These limitations reduce the resolution and detail in affected expression profiles and, consequently, restrict the amount of variance available for clustering genes algorithmically. RNA-seq datasets do not suffer from the plateau effect, but the RNA-seq dataset analyzed in this study was limited in that it was only performed in duplicate and only during the growth phase of the *Tetrahymena* life cycle. Additionally, despite the samples being synchronized, many gene expression observations were not repeated in the duplicated G1 and S phases, such as in Figure 4D. This was likely due to cell cycle synchronization diminishing over time. Overall, the tool would benefit from a new RNA-seq dataset that covers both the growth and conjugation phases with highly replicated samples collected at smaller time intervals. This would enhance both the precision and accuracy of the expression profiles and clustering algorithm’s ability to accurately partition genes. Naturally, more replicates in all conditions would further help to reduce the noise and improve clustering.

We highlighted only four biological functions in this report (mucocyst biogenesis and histone, ribosome, or proteasome processes), each with up to six associated co-expression clusters. However, our analysis produced hundreds of co-expression clusters that are enriched for biological functions (Table 2). These include metabolism, membrane trafficking, cytoskeletal organization, DNA replication, and many others (Supplementary File 3). A detailed analysis of all these gene modules is outside the scope of the present work, but it suggests the opportunity to target the study of many genes of interest in *Tetrahymena thermophila*.

One exciting extension of approaches like the TGNE will lie in their ability to elucidate cell biological pathways that evolved in specific lineages. As an example, we have previously found that co-expression analysis in *T. thermophila* could be leveraged to identify genes involved in a secretory protein complex that appears unique to the *Alveolata* lineage, which includes *Tetrahymena* and the apicomplexan *Toxoplasma gondii* (20). This indicates that signatures of functional association, as evidenced by co-expression patterns, persist and can therefore be informative through evolutionary time and speciation. In future work, tools like the TGNE for organisms that are chosen for their phylogenetic diversity, rather than for their experimental accessibility, could provide opportunities for translating experimental results between evolutionarily distant model systems, as well as for identifying lineage-specific cellular innovations.

## Author Contributions (CRediT taxonomy)

Conceptualization: LMZT, MAB, APT. Data Curation: LMZT, MAB, FY. Formal Analysis: LMZT, MAB. Funding Acquisition: APT, SK, SG, LMZT. Investigation: LMZT, MAB, LJB, YYJ, SK, DS, AP, FY. Methodology: LMZT, MAB, APT. Project Administration: LMZT, APT. Resources: APT, SG. Software: LMZT, MAB. Supervision: APT, LMZT. Validation: LMZT, MAB, APT. Visualization: LMZT, MAB. Writing - original draft: LMZT, MAB, APT. Writing - review and editing: LMZT, APT, MAB, FY, SG, SK, AP, DS

## Acknowledgments

We thank Daniela Sparvoli and Liam Elliot for their feedback on our manuscript and the TGNE. The University of Chicago Research Computing Center provided high performance computing resources and technical support. We are grateful to Benilton Carvalho (creator of the R oligo package) for technical support about RMA normalization in oligo. Wei Miao and Geoffrey Kapler released their *Tetrahymena* gene expression datasets publicly, and we appreciate the opportunity to analyze them. We thank Wei Miao for sharing negative control probe data for their original microarray study. Stijn van Dongen provided helpful discussions about co-expression clustering analysis. LMZT thanks the Stanford Energy Postdoctoral Fellowship, facilitated by the Precourt Institute for Energy at Stanford University, for supporting travel and publication costs for this work. APT’s laboratory was supported by NIH GM077607 and NSF MCB 1937326. LJB’s work was supported by NIH Genetics and Regulation Training Grant T32 GM007197. SK’s laboratory is funded by the Department of Biotechnology, India (BT/PR38584/MED/122/247/202), Department of Science and Technology (DST), India (CRG/2021/000732) and DBT/Wellcome Trust India Alliance (IA/I/22/2/506480). SG’s laboratory is supported by the National Natural Science Foundation of China (32125006 and 32070437).

## Supplementary Figure Captions

**Supplementary Figure 1.** Quality control of the microarray co-expression dataset. (A) Effect of RMA normalization on expression intensity distributions of all the microarray chips. Top: probability density of raw log2 expression intensities for each chip. Bottom: probability density of RMA normalized log2 expression intensities for each chip. (B) Box-and-whisker plots for the normalized unscaled standard error (NUSE) score of each chip. If NUSE = 1 is below the 25th percentile, the chip may be faulty. (C) A pseudo-image of a representative faulty chip that was identified by a high NUSE score, confirmed visually, and subsequently removed from the analysis. This chip, one of the replicates for the 12th hour starvation shows evidence of physical warping. Each probe is colored according to its rank of intensity with blue being lowest and red being highest. There should be no autocorrelation in the chip. (D) A hierarchical clustering of all chips that passed the quality control from (B) and (C). Two pairs of chips S0_GSM647651/S0_GSM647652 and S9_GSM647653/S9_GSM647654 are more similar to each other than to the other replicates for their respective experimental conditions. This is evidence of a batch effect, and this is supported by the fact that these chips were the only ones collected by a specific individual (Xiong et al., 2011). These four chips were removed from subsequent analysis. (E) The coefficient of variation versus the geometric mean of log2 expression for each gene in the final microarray dataset. Top: the unfiltered genes. Bottom: the genes that passed the expression filters. These filtered genes are the ones that were used for all subsequent analysis.

**Supplementary Figure 2.** Quality control of the RNA-seq co-expression dataset. (A) The relationship between the average Jaccard similarity of replicates and the CPM filtering threshold. This filter removes any gene that does not have a maximum CPM value above the threshold. The maximum Jaccard similarity is indicated by the red line, corresponding to a threshold of 0.802. (B) The effect of the threshold filter on the CPM distributions for every gene in every replicate condition. Genes that are retained are in blue, and genes that are filtered out are in red. The rightmost column displays the maximum CPM values for each gene across all replicates. (C) The average TPM values for each gene relative to its coefficient of variation before Jaccard filtering. (D) The average TPM values for each gene relative to its coefficient of variation after Jaccard filtering. Effectively, genes with a very low expression and a very high coefficient of variation are removed.

**Supplementary Figure 3.** 3D clustering parameter optimization plots for the Manhattan distance metric. (A) Min-max normalized microarray dataset. (B) Z-score normalized microarray dataset. (C) Min-max normalized RNA-seq dataset. (D) Z-score normalized RNA-seq dataset. We optimized for maximal modularity, maximal fraction of clusters with enriched functional terms, and minimal interquartile range for cluster size. Each curve corresponds to a different number of nearest neighbors, and each x along the curve scans across the Leiden clustering resolution parameter. Here, we are showing only the clustering based on Manhattan distance. The optimized partitions are circled in green, each corresponding to using three nearest neighbors and a resolution parameter of r = 0.005.

**Supplementary Figure 4.** 3D clustering parameter optimization plots for the Euclidean distance metric. (A) Min-max normalized microarray dataset. (B) Z-score normalized microarray dataset. (C) Min-max normalized RNA-seq dataset. (D) Z-score normalized RNA-seq dataset. We optimized for maximal modularity, maximal fraction of clusters with enriched functional terms, and minimal interquartile range for cluster size. Each curve corresponds to a different number of nearest neighbors, and each x along the curve scans across the Leiden clustering resolution parameter. Here, we are showing only the clustering based on Manhattan distance. The optimized partitions are circled in green, each corresponding to using three nearest neighbors and a resolution parameter of r = 0.005.

**Supplementary Figure 5.** 3D clustering parameter optimization plots for the CLR distance metric. (A) Min-max normalized microarray dataset. (B) Z-score normalized microarray dataset. (C) Min-max normalized RNA-seq dataset. (D) Z-score normalized RNA-seq dataset. We optimized for maximal modularity, maximal fraction of clusters with enriched functional terms, and minimal interquartile range for cluster size. Each curve corresponds to a different number of nearest neighbors, and each x along the curve scans across the Leiden clustering resolution parameter. Here, we are showing only the clustering based on Manhattan distance. The optimized partitions are circled in green, each corresponding to using four nearest neighbors and a resolution parameter of r = 0.005.

**Supplementary Figure 6.** 3D clustering parameter optimization plots for the angular distance metric. (A) Min-max normalized microarray dataset. (B) Z-score normalized microarray dataset. (C) Min-max normalized RNA-seq dataset. (D) Z-score normalized RNA-seq dataset. We optimized for maximal modularity, maximal fraction of clusters with enriched functional terms, and minimal interquartile range for cluster size. Each curve corresponds to a different number of nearest neighbors, and each x along the curve scans across the Leiden clustering resolution parameter. Here, we are showing only the clustering based on Manhattan distance. The optimized partitions are circled in green, each corresponding to using three nearest neighbors and a resolution parameter of r = 0.005.

**Supplementary Figure 7.** 3D clustering parameter optimization plots for the linear correlation distance metric. (A) Min-max normalized microarray dataset. (B) Z-score normalized microarray dataset. (C) Min-max normalized RNA-seq dataset. (D) Z-score normalized RNA-seq dataset. We optimized for maximal modularity, maximal fraction of clusters with enriched functional terms, and minimal interquartile range for cluster size. Each curve corresponds to a different number of nearest neighbors, and each x along the curve scans across the Leiden clustering resolution parameter. Here, we are showing only the clustering based on Manhattan distance. The optimized partitions are circled in green, each corresponding to using three nearest neighbors and a resolution parameter of r = 0.005.

**Supplementary Figure 8.** Quality control of mucocyst replacement experiment. (A) Pseudo-images of the microarray chips, colored by rank of probe intensity (red is high, blue is low). (B) Box-and-whisker plots of the normalized unscaled standard error (NUSE) for each chip. The seventh chip was removed from the subsequent analysis because its 25% percentile for the NUSE score was significantly above 1.

**Supplementary Figure 9.** Enrichment, differential expression, and overlap of experimentally validated mucocyst-associated and differentially expressed, upregulated genes. Z-score normalized expression profiles for genes in (A) the four microarray and (B) the four RNA-seq clusters significantly enriched for experimentally validated mucocyst-associated genes as well as the 33 genes overlapping between the upregulated, enriched microarray clusters, and enriched RNA-seq clusters in (C) the microarray and (D) the RNA-seq datasets. (E) Volcano plot illustrating differential expression of each gene represented in the microarray dataset over one hour in the MN173 mutant relative to the wild type *T. thermophila*. This is the same plot as panel Figure 2E. Thresholds are represented by blue dashed lines (q < 0.01 and fold-change > 1.5). All genes that passed the thresholds have a Bayesian posterior probability of differential expression greater than 80%. (F) Venn diagram describing the overlapping genes in the enriched microarray clusters, enriched RNA-seq clusters, and the set of upregulated genes with min-max normalization. Min-max normalized expression profiles for genes that are co-expressed in the microarray and RNA-seq datasets, but not detected in the upregulated dataset: (G) gene expression in the microarray profiles and (H) gene expression in the RNA-seq profiles.

**Supplementary Figure 10.** Comparison of GRL expression profiles and clustering. (A) Z-score normalized GRL expression profiles in the microarray dataset. There are two major clusters–– GRL3, 5, 7, 8 and GRL1, 2, 4––and GRL6 and GRL9 do not conform to either one. The distinction between the two major clusters indicates that the cluster containing GRL3, 5, 7, 8 begins to lose expression one time point later during the starvation and conjugation conditions than does the one containing GRL1, 2, 4. (B) Z-score normalized GRL expression profiles in the RNA-seq dataset. During the mitotic cell cycle, GRL1, 3, 4, 5, 7, 8 cluster together and GRL2, 9 cluster together. GRL6 presents a very different expression profile from the other GRLs.

