## Supplementary Figure 1 for "Inferring gene-pathway associations from consolidated transcriptome datasets: an interactive gene network explorer for *Tetrahymena thermophila*"

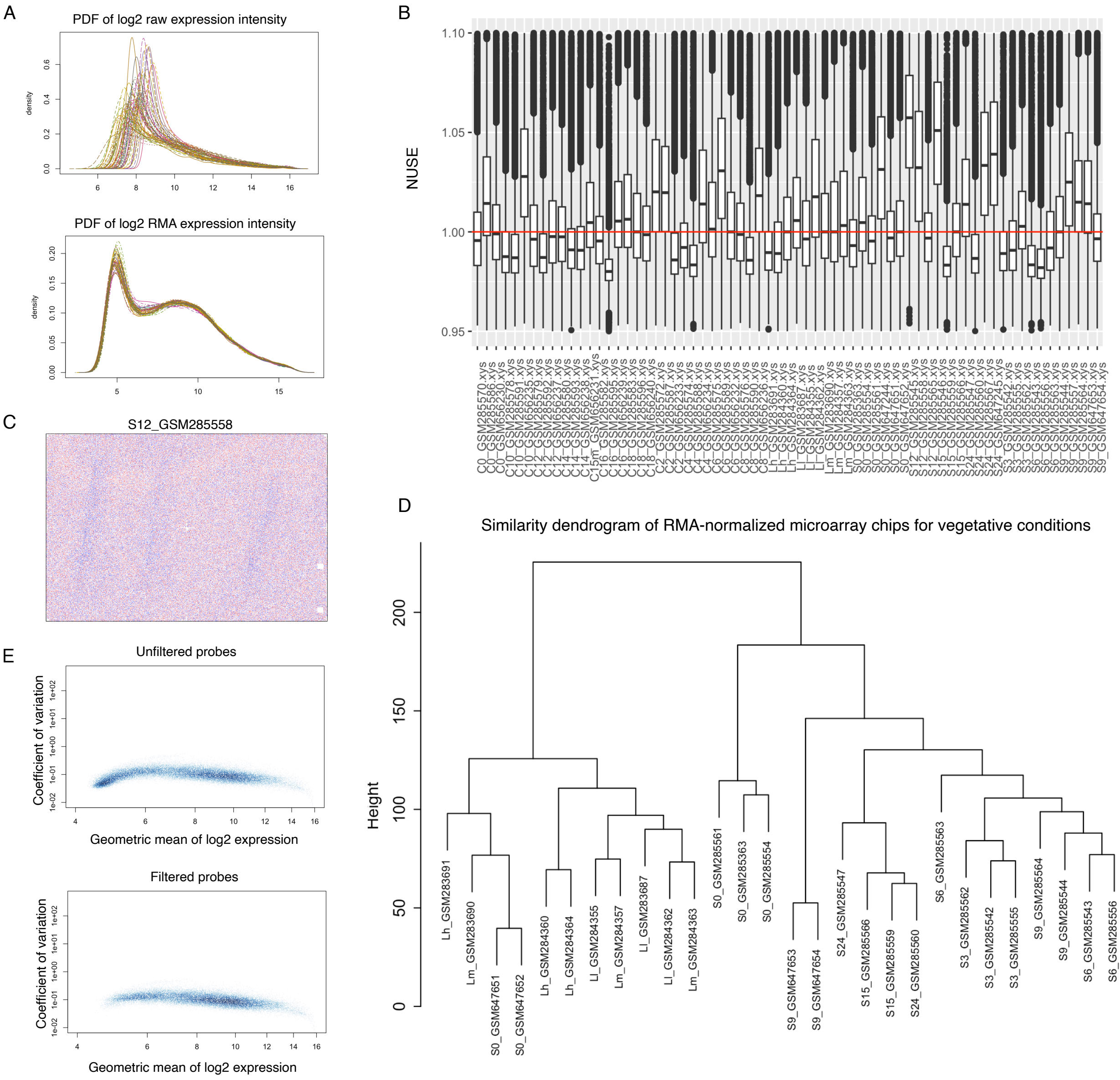


**Supplementary Figure 1.** Quality control of the microarray co-expression dataset. (A) Effect of RMA normalization on expression intensity distributions of all the microarray chips. Top: probability density of raw log2 expression intensities for each chip. Bottom: probability density of RMA normalized log2 expression intensities for each chip. (B) Box-and-whisker plots for the normalized unscaled standard error (NUSE) score of each chip. If NUSE = 1 is below the 25th percentile, the chip may be faulty. (C) A pseudo-image of a representative faulty chip that was identified by a high NUSE score, confirmed visually, and subsequently removed from the analysis. This chip, one of the replicates for the 12th hour starvation shows evidence of physical warping. Each probe is colored according to its rank of intensity with blue being lowest and red being highest. There should be no autocorrelation in the chip. (D) A hierarchical clustering of all chips that passed the quality control from (B) and (C). Two pairs of chips S0_GSM647651/S0_GSM647652 and S9_GSM647653/S9_GSM647654 are more similar to each other than to the other replicates for their respective experimental conditions. This is evidence of a batch effect, and this is supported by the fact that these chips were the only ones collected by a specific individual (Xiong et al., 2011). These four chips were removed from subsequent analysis. (E) The coefficient of variation versus the geometric mean of log2 expression for each gene in the final microarray dataset. Top: the unfiltered genes. Bottom: the genes that passed the expression filters. These filtered genes are the ones that were used for all subsequent analysis.
