## Supplementary Figure 2 for "Inferring gene-pathway associations from consolidated transcriptome datasets: an interactive gene network explorer for *Tetrahymena thermophila*"

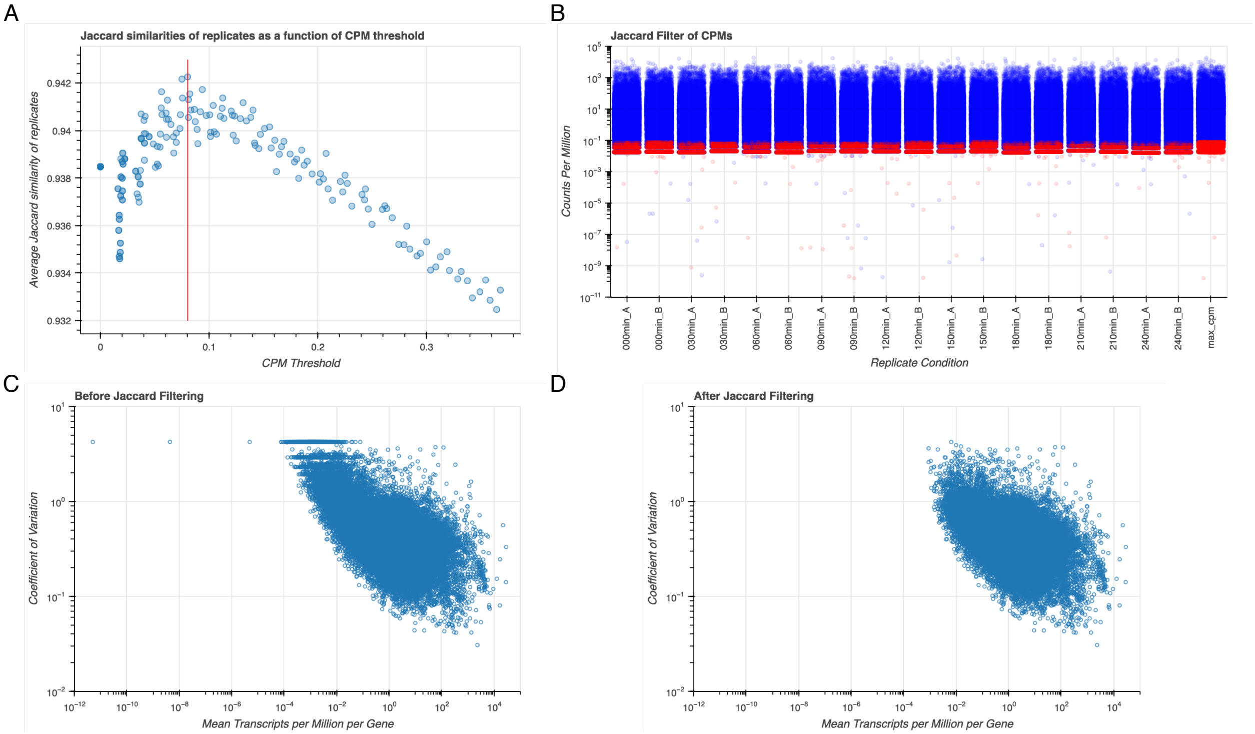


**Supplementary Figure 2.** Quality control of the RNA-seq co-expression dataset. (A) The relationship between the average Jaccard similarity of replicates and the CPM filtering threshold. This filter removes any gene that does not have a maximum CPM value above the threshold. The maximum Jaccard similarity is indicated by the red line, corresponding to a threshold of 0.802. (B) The effect of the threshold filter on the CPM distributions for every gene in every replicate condition. Genes that are retained are in blue, and genes that are filtered out are in red. The rightmost column displays the maximum CPM values for each gene across all replicates. (C) The average TPM values for each gene relative to its coefficient of variation before Jaccard filtering. (D) The average TPM values for each gene relative to its coefficient of variation after Jaccard filtering. Effectively, genes with a very low expression and a very high coefficient of variation are removed.
