## Supplementary Figure 3 for "Inferring gene-pathway associations from consolidated transcriptome datasets: an interactive gene network explorer for *Tetrahymena thermophila*"

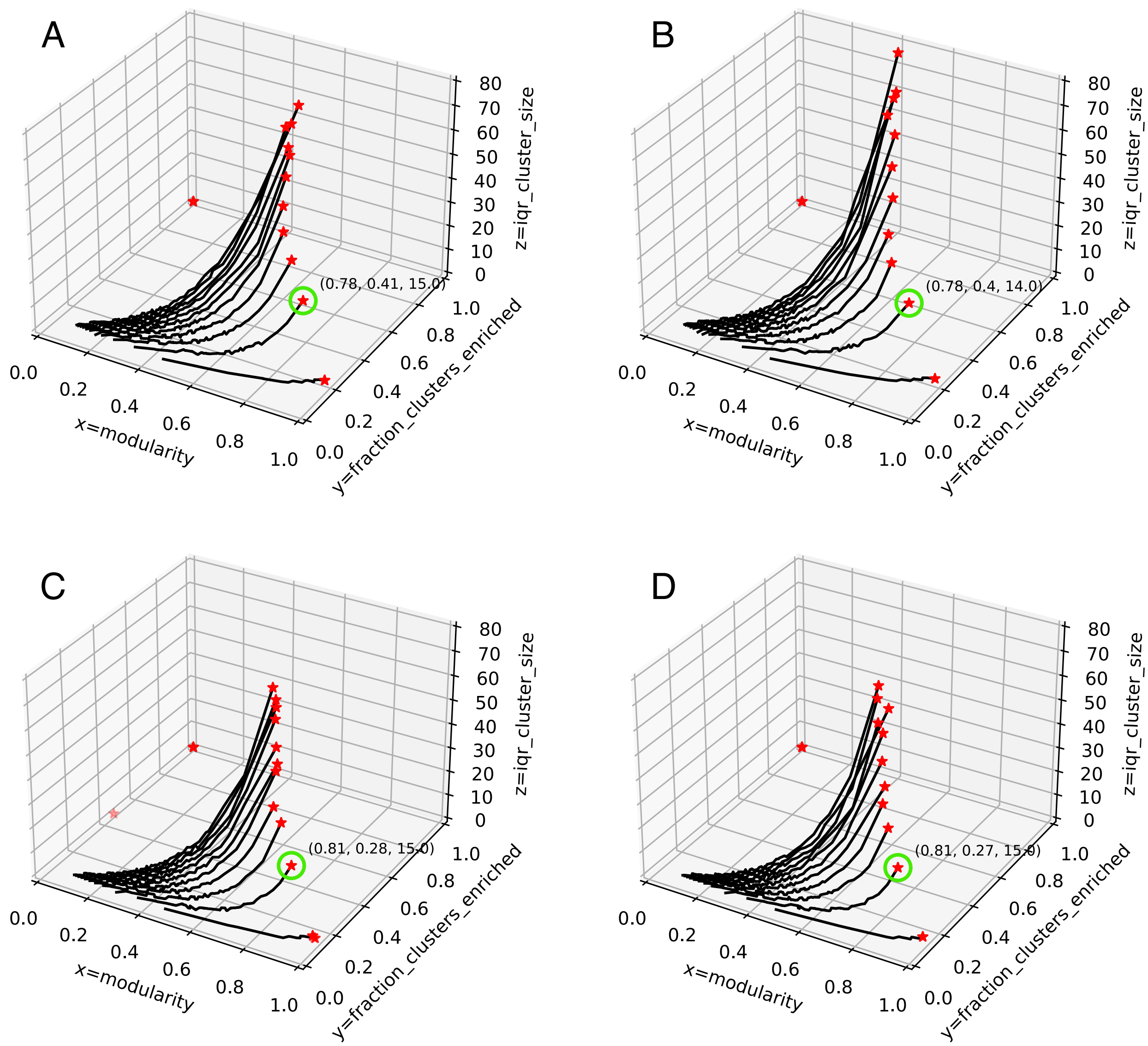


**Supplementary Figure 3.** 3D clustering parameter optimization plots for the Manhattan distance metric. (A) Min-max normalized microarray dataset. (B) Z-score normalized microarray dataset. (C) Min-max normalized RNA-seq dataset. (D) Z-score normalized RNA-seq dataset. We optimized for maximal modularity, maximal fraction of clusters with enriched functional terms, and minimal interquartile range for cluster size. Each curve corresponds to a different number of nearest neighbors, and each x along the curve scans across the Leiden clustering resolution parameter. Here, we are showing only the clustering based on Manhattan distance. The optimized partitions are circled in green, each corresponding to using three nearest neighbors and a resolution parameter of r = 0.005.
