## Supplementary Figure 8 for "Inferring gene-pathway associations from consolidated transcriptome datasets: an interactive gene network explorer for *Tetrahymena thermophila*"

A

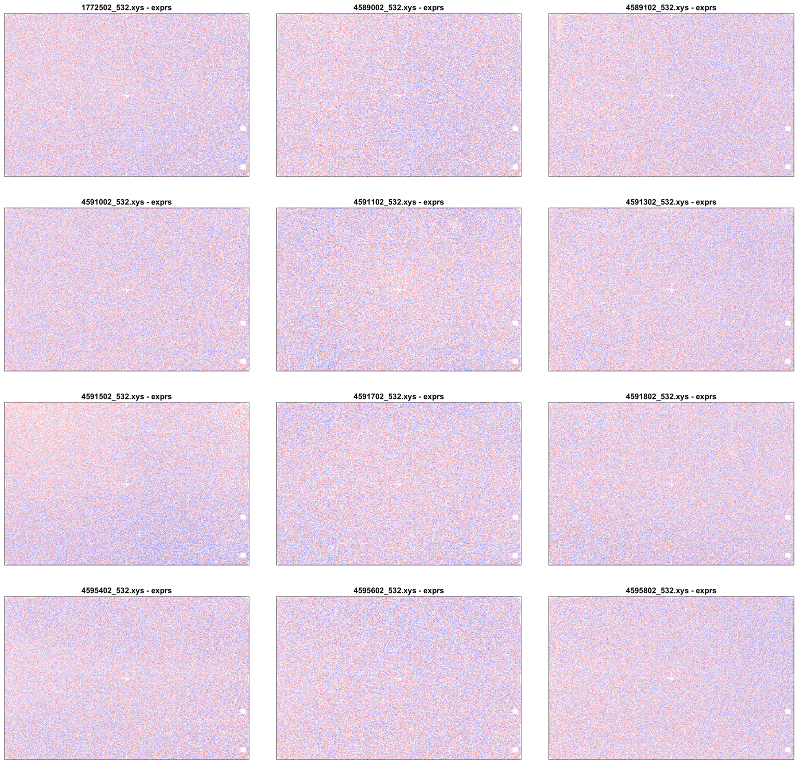


B

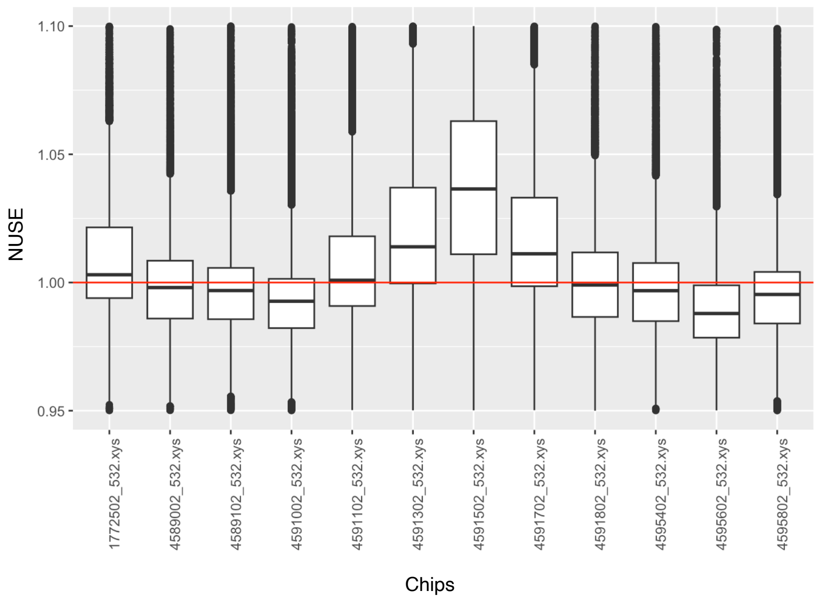


**Supplementary Figure 8.** Quality control of mucocyst replacement experiment. (A) Pseudo-images of the microarray chips, colored by rank of probe intensity (red is high, blue is low). (B) Box-and-whisker plots of the normalized unscaled standard error (NUSE) for each chip. The seventh chip was removed from the subsequent analysis because its 25% percentile for the NUSE score was significantly above 1.
