## Supplementary Figure 9 for "Inferring gene-pathway associations from consolidated transcriptome datasets: an interactive gene network explorer for *Tetrahymena thermophila*"

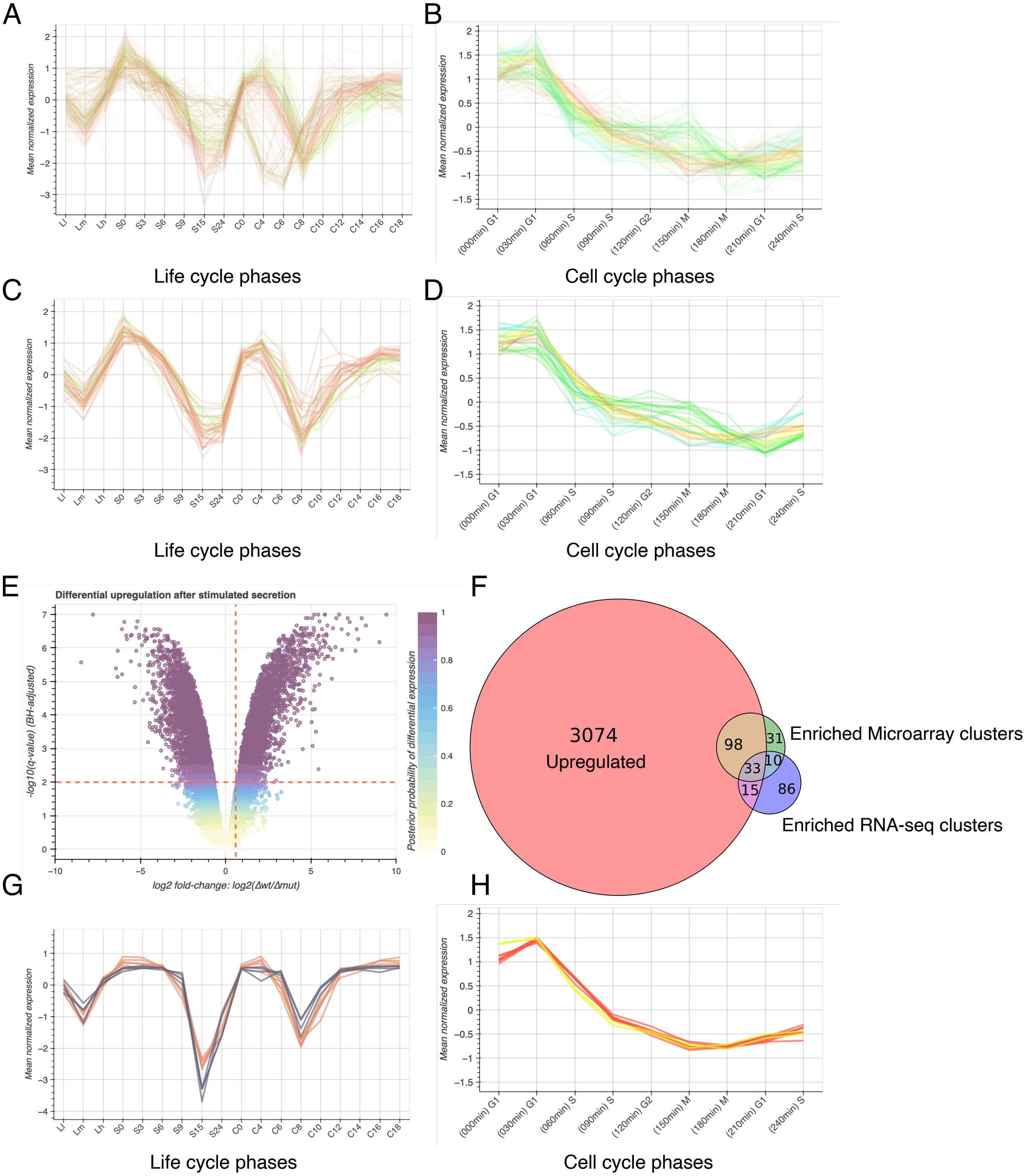


**Supplementary Figure 9.** Enrichment, differential expression, and overlap of experimentally validated mucocyst-associated and differentially expressed, upregulated genes. Z-score normalized expression profiles for genes in (A) the four microarray and (B) the four RNA-seq clusters significantly enriched for experimentally validated mucocyst-associated genes as well as the 33 genes overlapping between the upregulated, enriched microarray clusters, and enriched RNA-seq clusters in (C) the microarray and (D) the RNA-seq datasets. (E) Volcano plot illustrating differential expression of each gene represented in the microarray dataset over one hour in the MN173 mutant relative to the wild type *T. thermophila*. This is the same plot as panel Figure 2E. Thresholds are represented by blue dashed lines ($q < 0.01$ and fold-change > 1.5). All genes that passed the thresholds have a Bayesian posterior probability of differential expression greater than 80%. (F) Venn diagram describing the overlapping genes in the enriched microarray clusters, enriched RNA-seq clusters, and the set of upregulated genes with min-max normalization. Min-max normalized expression profiles for genes that are co-expressed in the microarray and RNA-seq datasets, but not detected in the upregulated dataset: (G) gene expression in the microarray profiles and (H) gene expression in the RNA-seq profiles.
