## Supplementary Figure 10 for "Inferring gene-pathway associations from consolidated transcriptome datasets: an interactive gene network explorer for *Tetrahymena thermophila*"

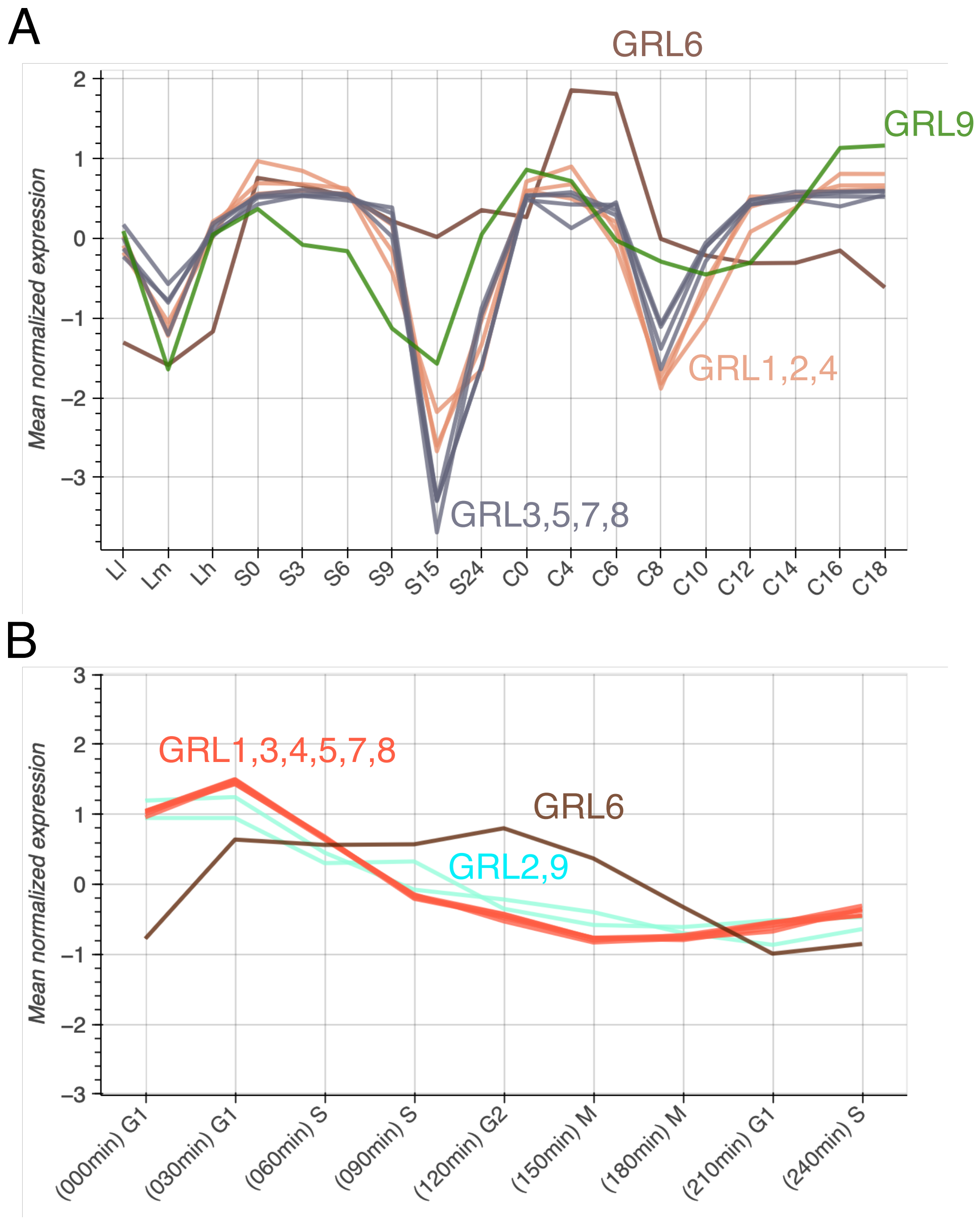


**Supplementary Figure 10.** Comparison of GRL expression profiles and clustering. (A) Z-score normalized GRL expression profiles in the microarray dataset. There are two major clusters—GRL3,5,7,8 and GRL1,2,4—and GRL6 and GRL9 do not conform to either one. The distinction between the two major clusters indicates that the cluster containing GRL3,5,7,8 begins to lose expression one time point later during the starvation and conjugation conditions than does the one containing GRL1,2,4. (B) Z-score normalized GRL expression profiles in the RNA-seq dataset. During the mitotic cell cycle, GRL1,3,4,5,7,8 cluster together and GRL2,9 cluster together. GRL6 presents a very different expression profile from the other GRLs.
